# UCHL1 facilitates aggregates clearance and enhances neural stem cell activation in spinal cord injury

**DOI:** 10.1101/2021.09.23.461600

**Authors:** Lu Ding, Weiwei Chu, Yu Xia, Ming Shi, Tian Li, Feng-Quan Zhou, David Y.B. Deng

**Affiliations:** Scientific Research Center, The Seventh Affiliated Hospital, Sun Yat-sen University, Shenzhen 518107, China; Obstetrics and Gynecology Department, The Seventh Affiliated Hospital, Sun Yat-sen University, Shenzhen 518107, China; School of Pharmaceutical Sciences (Shenzhen), Sun Yat-sen University, Shenzhen 518107, China; Department of Orthopaedic Surgery and Department of Neuroscience, Johns Hopkins University School of Medicine, Baltimore, Maryland 21287, USA; Sir Run Run Shaw Hospital, Zhejiang University School of Medicine, Hangzhou 310016, China

**Keywords:** Ubiquitin c-terminal hydrolase l-1, protein aggregates clearance, neural stem cell activation, reactive astrocyte, complement component 3, proteasome, spinal cord injury

## Abstract

Activation of endogenous neural stem cells (NSCs) is critically important for the adult neurogenesis. However, NSC activation is extremely limited after spinal cord injury (SCI). Recent evidence suggests that accumulation of protein aggregates impedes quiescent NSC activation. Here, we found ubiquitin c-terminal hydrolase l-1 (UCHL1), an important deubiquitinating enzyme, functioned to facilitate NSC activation by clearing protein aggregations through ubiquitin-proteasome approach. Upregulation of UCHL1 enhanced NSC proliferation in the spinal cord after injury. Based on protein microarray analysis of SCI cerebrospinal fluid, it is further revealed that C3^+^ neurotoxic reactive astrocytes negatively regulated UCHL1 and aggresome clearance through C3/C3aR signaling, resulting in reduced capacity of NSC to activate. Furthermore, blockade of reactive astrocytes or C3/C3aR pathway led to enhanced NSC activation post-SCI. Together, this study elucidated a mechanism regulating NSC activation in the adult spinal cord involving the UCHL1-proteasome approach, which may provide potential molecular targets for NSC fate regulation.

## INTRODUCTION

Brain and spinal cord injuries typically results in permanent neurological impairment with no curative treatment available. Endogenous neural stem cells (NSCs) present in the adult central nervous system (CNS) could be an attractive candidate for CNS repair as an alternative to stem cell transplantation therapy. Despite promising progress of NSCs in neurogenesis in the forebrain, little is known concerning NSCs in the spinal cord, a non-neurogenic region. Despite that ependymal cells have long been identified as the resident NSCs in the adult spinal cord (Barnabe-Heider et al., 2010; Meletis et al., 2008), it is currently a great controversial issue that which type of cells are the resident spinal cord NSCs for the contradictory conclusions concerning their stemness in several researches (Barnabe-Heider *et al*., 2010; Garcia-Ovejero et al., 2015; Muthusamy et al., 2018; Ren et al., 2017; Shah et al., 2018). Nestin, known as a neuroepithelial stem cell protein, is used as a good marker for neural stem/progenitors cells of CNS in numerous studies (Bernal and Arranz, 2018; Morrow et al., 2020; Tobin et al., 2019). Nestin^+^ cells from the spinal cord could form neurospheres and differentiate into neurons, astrocytes and oligodendrocytes in vitro (Barnabe-Heider *et al*., 2010; Meletis *et al*., 2008). Recent single-cell RNA sequencing analysis also revealed that genes of active NSCs were significantly upregulated in Nestin^+^ cells and they exhibited the differentiated intermediate states after SCI, suggesting that Nestin^+^ cells may act as the endogenous NSCs in the spinal cord (Shu et al., 2021). Nestin^+^ NSCs are quiescent in the normal spial cord and remarkably increased after spinal cord injury (SCI)(Shu *et al*., 2021). Unfortunately, although quickly activated in response to SCI, the activation and neurogenesis from endogenous NSCs are extremely limited in the adult spinal cord in vivo (Barnabé-Heider et al., 2010; Liu et al., 2015; McTigue and Sahinkaya, 2011). Such non-neurogenic feature of injured spinal cord suggests the presence of negative influences from the spinal cord microenvironment on NSCs. Therefore, elucidating the potential mechanisms by which spinal cord microenvironmental and intrinsic factors regulate the endogenous NSC activation and neurogenesis is of great importance for SCI repair.

Maintenance of protein homeostasis (proteostasis) is vital for proper cell function. As a result, imbalance of proteostasis often results in accumulation of misfolded and abnormal proteins, which is closely associated with stem cell dysfunction, aging and neurodegenerative disorders (Vilchez et al., 2014). The ubiquitin-proteasome system (UPS) and the lysosome/autophagy proteolytic system are the primary mechanisms responsible for maintaining protein homeostasis (Samant et al., 2018; Wong and Cuervo, 2010). Various proteins and regulators involved in mammalian neurogenesis and NSC physiology are directly proteolytic degradation by UPS, suggesting the critical role of UPS in adult neurogenesis and NSC regulation (Naujokat, 2009a). Recent transcriptomic data (Leeman et al., 2018) showed that the clearance of protein aggregates mediated by lysosomes/proteasome systems plays critical roles in the regulation of NSC activation. Specifically, the quiescent NSCs in aged mouse brains have reduced activation capacity due to lysosome defects and accumulation of protein aggregates, whereas enhancing lysosome function restored their ability to be activated (Leeman *et al*., 2018). In addition, Morrow et al. (Morrow *et al*., 2020) found that vimentin could recruit the aggresome to the proteasome machinery to clear aberrant proteins during adult quiescent NSCs activation. Conversely, knocking out vimentin in NSCs impaired proteostasis recovery and delayed its quiescence exit, suggesting the crucial effects of UPS in NSC activation.

Ubiquitin C-terminal Hydrolase L1 (UCHL1), a member of deubiquitinating enzymes abundantly expressed in central nervous system (CNS), is indispensable for the proper UPS functions. It hydrolyzes free mono-ubiquitin from ubiquitinated proteins for recycling, ligate ubiquitin (ub) into specific proteins for proteasome degradation and also can bind to free monoubiquitin to maintain an available ubiquitin pool (Cartier et al., 2009; Wilkinson et al., 1989). As an important player in abnormal protein aggregates clearance, UCHL1 plays critical role in several neurodegenerative diseases and CNS trauma (Bilguvar et al., 2013; Gong et al., 2006; Graham and Liu, 2017; Liu et al., 2002). For instance, the protein level of UCHL1 is inversely associated with the number of neurofibrillary tangles in AD brains (Choi et al., 2004); the overexpression of it resulted in decreased level of phosphorylated tau, while its knockdown led to increased formation of phosphorylated tau (Cartier *et al*., 2009). It also preserves axonal conductance and improves motor behavior after stroke via decreases polyubiquitinated protein accumulation (Liu et al., 2019). In addition, research has reported that UCHL1 could spatially mediates neurogenesis in the embryonic brain by regulating the morphology of neural progenitor cells (NPCs) (Sakurai et al., 2006). However, the potential effects of UCHL1 on spinal cord Nestin^+^ NSCs remain unclear. Given the pivotal role of UCHL1 in removing protein aggregations, it may coordinate UPS functions in controlling endogenous NSC activation.

The pathological milieu of spinal cord is a vital factor controlling proteasome functions, which likely underlies the limited activation of endogenous NSCs and their non-neuronal cell fate determination post-SCI. As the most abundant glia cells in CNS, astrocytes play important roles in the creation of neurogenic micro-environment (Freeman, 2010; Karimi-Abdolrezaee and Billakanti, 2012). In response to CNS diseases or injuries, astrocytes usually undergo inflammatory transitions, resulting in activation of immune mediators and the release of proinflammatory cytokines (Escartin et al., 2021; Liddelow and Barres, 2017). Reactive astrocytes have been reported to mediate cellular toxicity to neurons and oligodendrocytes by secretion of complement component 3 (C3) and lipoparticle proteins such as Apolipoprotein E (APOE) and Apolipoprotein J (APOJ) (Guttenplan et al., 2021). Moreover, C3 has also been implicated in the accumulation of neurotoxic proteins within neurons in multiple neurodegeneration disorders (Hong et al., 2016; Liddelow et al., 2017; Wu et al., 2019). However, the impact of neurotoxic reactive astrocytes on UCHL1-proteasome functions and NSC activation is unknown.

Here, we explored the potential effects of neurotoxic reactive astrocytes and UCHL1-mediated protein aggregates clearance on spinal Nestin^+^ NSC activation following SCI. The present study showed that the level of UCHL1 decreased after SCI and its upregulation facilitated Nestin^+^ NSC activation by eliminating protein aggregations through ubiquitin-proteasome pathway (UPP). Based on protein microarray analysis of cerebrospinal fluid (CSF), we found that C3 and the complement pathway were remarkably upregulated post-SCI, and it further revealed that neurotoxic reactive astrocytes induced by SCI inhibited UCHL1 and UPP through C3 secretion, resulting in protein aggregates accumulation in NSCs and their decreased ability to activate. Blockade of reactive astrocytes or C3/C3aR pathway both promoted Nestin^+^ NSC proliferation in the adult spinal cord after SCI. Together, this study suggested that SCI-induced reactive astrocytes could hinder the activation of endogenous Nestin/SOX2^+^ NSCs by repressing the UCHL1-UPP via C3/C3aR signaling, thus providing potential therapeutic targets for the stem cell-based therapies of CNS disorders.

## RESULTS

### The expression of UCHL1 in the spinal cord was significantly decreased after SCI

Trauma to CNS generally cause dysfunctions of UPP, resulting in depletion of free ubiquitin and accumulation of ubiquitinated proteins. Therefore, in this study we investigated several isoforms of UCHL (UCHL1, UCHL3, UCHL5) in the spinal cord that act to maintain free ubiquitin level and proper UPS function. Results showed that UCHL1, a neuron-specific isoform, was strongly detected in the brain and the spinal cord, compared with the other two isoforms UCHL3 and UCHL5 (Figure 1, A-C). At 24 hours after SCI, a remarkable down-regulation of UCHL1 level was observed in the gray matter (Figure 1, D and E), and it remained to be significantly decreased within three days post-injury (Figure 1, F and G). Further immunofluorescent staining showed that UCHL1 positive cells in the spinal cord did not overlap with NG2^+^ oligodendrocyte precursor cells (OPCs) or GFAP^+^ astrocytes (Figure 1H). Instead, they were highly enriched in the spinal NeuN^+^ neurons and Nestin^+^ cells, a specific marker for NSCs (Hu et al., 2019; Kim et al., 2019; Shu *et al*., 2021) (Figure 1H). More importantly, the expression of UCHL1 was remarkably decreased in both of neurons and Nestin^+^ NSCs after SCI (Figure 1, H and I), suggesting that it might be involved in regulation of spinal cord neurons and NSCs function.

**Figure 1.**
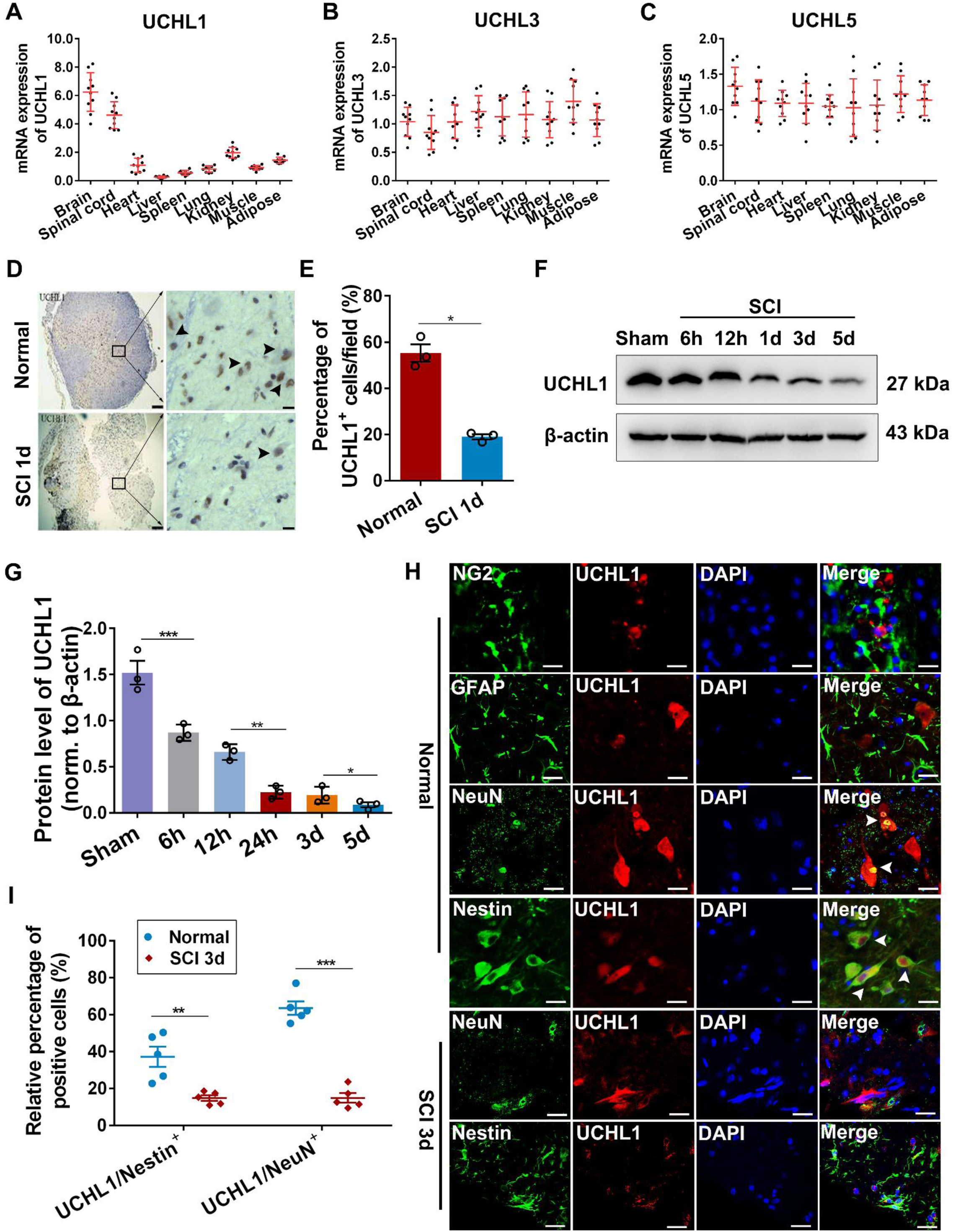
UCHL1 was expressed in the spinal Nestin^+^ NSCs and downregulated remarkably after SCI. (A-C) QPCR analysis showing the relative expression of UCHL1, UCHL3 and UCHL5 in the different organs of rats. UCHL1 was abundantly expressed in the brain and spinal tissues of rats. n=10 independent animals. (D-E) Immunohistochemical representative images (D) and quantification (E) showed that UCHL1 was remarkably downregulated in the gray matter at 24 h after SCI. Scale bar (left), 100 μm, Scale bar (right), 50μm. (E) n=3 independent animals. Data are presented as mean ± SEM. p-values (*p<0.05) are calculated using two-tailed unpaired Student’s t-test. (F-G) Western blot analysis showed the protein level of UCHL1 in the spinal cord post-SCI. Representative immunoblot images and quantification of Western blot assay was shown in F and G. (G) n=3 independent animals. (H) Immunofluorescence assay shows that UCHL1 was expressed within NeuN^+^ neurons and Nestin^+^ NSCs, but not NG2^+^ oligodendrocyte precursor cells and GFAP^+^ astrocytes. Scale bar, 25 μm. Data are presented as mean ± SEM. p-values (*p<0.05, **p<0.01, ***p<0.001) are calculated using one-way ANOVA with Tukey HSD post hoc test. (I) The relative percentage of UCHL1/Nestin^+^ cells and UCHL1/ NeuN^+^ neurons before and at 3 days after SCI. n=5 independent animals. Data are presented as mean ± SEM. p-values (**p<0.01, ***p<0.001) are calculated using (E/I) two-tailed unpaired Student’s t-test.

### Upregulation of UCHL1 promoted NSC activation and proliferation by ubiquitin-proteasome approach-dependent protein aggregates clearance in vitro

Evidence is emerging that there is a significant difference of protein aggregates and UPS activity between the activated and quiescent NSCs (Leeman *et al*., 2018). We first explored whether UCHL1 was involved in the protein aggregates removal within NSCs. NSCs in the brain or spinal cord exhibit same capability of self-renewing and multipotency (Grégoire et al., 2015). Due to the limited number of endogenous NSCs in adult spinal cord, NSCs extracted from the brain of fetal rats were utilized. Cells were infected with either empty lentiviral vector (NC-LV), lentiviral vector encoding UCHL1 (OE-UCHL1-LV), or treated with LDN-57444, an effective competitive and site-directed enzyme activity inhibitor of UCHL1 (Gong *et al*., 2006). The successful overexpression of UCHL1 in NSCs was confirmed at 48 hours after infection (Figure 2, A-C). Then the intracellular aggregations were identified by Proteostat dye. Compared with many protein aggregates accumulated in the control NSCs, those infected with OE-UCHL1-LV exhibited less Proteostat staining (Figure 2D-F and Supplementary Figure1 A-B), while LDN-57444 treatment led to much more protein aggregates (Figure 2, D-F and Supplementary Figure1, A and B). No difference was observed between the control (no viral vector infection) and NC-LV group, indicating that lentiviral vector *per se* had no effect.

**Figure 2.**
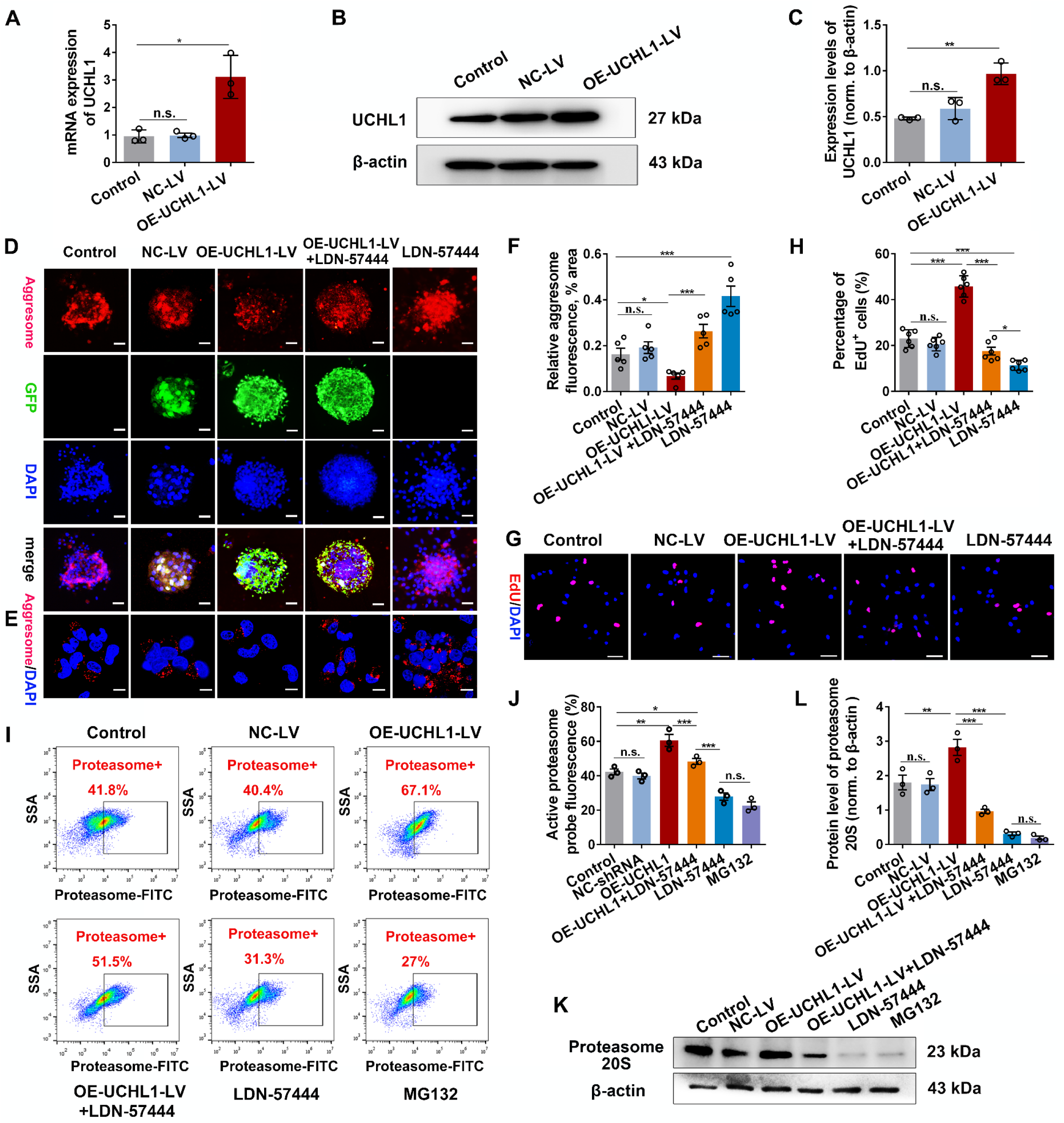
Overexpression of UCHL1 promoted NSC proliferation by accelerating protein aggregates clearance via ubiquitin-proteasome pathway. (A) QPCR analysis showing the mRNA level of UCHL1 in NSCs transfected with NC-LV or OE-UCHL1-LV for 48h. n=3 biological replicates. Data are presented as mean ± SEM. p-values (*p<0.05, n.s. not significant) are calculated using two-tailed unpaired Student’s t-test. (B-C) The protein expression of UCHL1 in NSCs transfected with OE-UCHL1-LV for 48h by Western blot analysis. Representative images and quantification of Western blot analysis were shown in B and C. (C) n=3 biological replicates. Data are presented as mean ± SEM. p-values (**p<0.01, n.s. not significant) are calculated using two-tailed unpaired Student’s t-test. (D-E) Confocal representative images showing the enrichment of protein aggregates (aggresome^+^) in NSCs transfected among different groups. Scale bar (D), 100 μm, Scale bar (E), 10 μm. (F) Quantification of protein aggregates (aggresome^+^) in NSCs transfected with NC-LV or OE-UCHL1-LV. n=5 biological replicates. Data are presented as mean ± SEM. p-values (**p<0.01, ***p<0.001, n.s. not significant) are calculated using Kruskal Wallis test with Bonferroni correction. (G-H) Confocal representative images and quantification of the proliferating NSCs (EdU/Nestin^+^) in different treatments. Scale bar (G), 10 μm. (H) n=6 biological replicates. (I) The proteasome activity of NSCs in groups was measured by flow cytometry assay. (J) Quantification of proteasome activity in NSCs by flow cytometry assay. n=3 biological replicates. (K-L) Western blot analysis was performed to detected the protein level of 20S proteasome in NSCs. Representative immunoblot images and quantification of relative protein level were shown in K and L. (L) n=3 biological replicates. (F/H/J/L) Data are presented as mean ± SEM. p-values (*p<0.05, **p<0.01, ***p<0.001, n.s. not significant) are calculated using one-way ANOVA with Tukey HSD post hoc test. See also supplementary Figure 1.

A recent study reported that increased protein aggregates accumulation in quiescent NSCs reduced their ability to be activated (Leeman *et al*., 2018). Thus, we examined whether UCHL1 regulated NSC proliferation using the EdU incorporation assay. Results showed that upregulation of UCHL1 (OE-UCHL1-LV) promoted NSCs proliferation, which was abolished by LDN-57444 (Figure 2, G and H; Supplementary Figure 1, C-D). Furthermore, up-regulation of UCHL1 in NSCs led to more Tubulin β3^+^ neurons, whereas NSCs treated with LDN-57444 mainly differentiated into astrocytes (Supplementary Figure 1, E-H). These evidences suggested that upregulation of UCHL1 may facilitate NSCs proliferation and neuronal differentiation by accelerating the clearance of intracellular protein aggregates in vitro.

To test this hypothesis, we first examined if manipulation of protein aggregates in NSCs could affect their activation. Because nutrient deprivation could significantly reduce protein aggregates in NSCs (Leeman *et al*., 2018), we incubated NSCs in HBSS (nutrient deprivation) prior to the treatment with activation factors epidermal growth factor (EGF) and fibroblast growth factor (FGF). The results showed that nutrient deprivation depleted protein aggregations in NSCs (Supplementary Figure 2, A-D) and enhanced NSC activation in response to growth factors (Supplementary Figure 2, E-H); in contrast, administration of MG132, a proteasome inhibitor, resulted in accumulation of protein aggregates (Supplementary Figure 2, A-D) and markedly impeded NSC proliferation (Supplementary Figure 2, E-H), implying protein aggregations in NSCs directly impacted its activation. We next determined if UCHL1 activated NSCs via UPP-dependent clearance of protein aggregates by monitoring the proteasome activity using the proteasome-specific affinity probe. Enhanced proteasome activity was detected in the OE-UCHL1-LV treatment group, while NSCs treated with LDN-57444 or MG132 led to significantly lower proteasome activities compared with the control (Figure 2, I and J). The protein level of proteasome 20S, the catalytic core of the 26S proteasome that act as the central protease of the ubiquitin pathway of protein degradation, was increased accordingly after UCHL1 upregulation (Figure 2, K and L). Collectively, these data provided evidence that UCHL1 could activate NSCs via the UPS-dependent clearance of protein aggresome.

### Reactive astrocytes suppressed NSC activation by inhibiting UCHL1 and protein aggregation elimination

How does SCI lead to reduced UCHL1 level and failed activation of endogenous NSCs? CSF directly contacts with the CNS and contains various biomarkers reflecting the damage to brain/spinal cord or neurodegenerative diseases. Thus, we used protein array analysis to compare a spectrum of cytokines in CSF before and after SCI. From the hierarchical clustering analysis, 52 of total 88 cytokines were significantly upregulated after SCI, among which 26 showed more than 2-fold changes (Supplementary Figure 3A). Consistent with previous studies (Benedet et al., 2021; Jha and Suk, 2013), the level of several astroglia-related factors, including GFAP, S100B, S100A, were all markedly increased in CSF of SCI rats (Supplementary Figure 3A), which might be indicators for astroglial activation. Interestingly, C3, the central component of classical complement pathway, was also increased significantly. KEGG analysis revealed that the significantly altered cytokines were mainly linked to signaling pathways associated with immune-inflammation responses, such as the complement and coagulation cascades, TNF and NFκB signaling pathways, etc. (Supplementary Figure 3B). The increased level of C3a was also detected in the CSF and serum of SCI rats (Supplementary Figure 3, C and D), and in the damaged spinal tissues (Supplementary Figure 3, E and F). It has been reported that C3 is one of the typical biomarkers of neurotoxic reactive astrocytes (Liddelow *et al*., 2017; Wu *et al*., 2019). Indeed, we found that C3/GFAP^+^ reactive astrocytes were significantly increased after SCI (Supplementary Figure 3, G and H). We thus proposed that acute insult to spinal cord may lead to activation of neurotoxic reactive astrocytes, which subsequently suppress UCHL1 and UPS functions in NSCs, resulting in the dysfunction of abnormal protein aggregates removal.

To test this idea, we prepared purified primary astrocytes deprived from fetal rats, and induced neurotoxic reactive phenotype with three cytokines TNFα/IL-1α/C1q, the best inducers of reactive astrocytes, as previously described (Guttenplan *et al*., 2021; Liddelow *et al*., 2017). The purity of astrocytes was more than 96% (Supplementary Figure 4, A-C). Double-staining of GFAP with microglia marker TMEM119 or neuronal marker NeuN further ruled out the potential contribution of other cell types, such as microglia, in the culture system (Supplementary Figure 4, D and E). Firstly, C3/GFAP^+^ reactive astrocytes were induced successfully (Figure 3, A-B). Next, a co-culture system was constructed to evaluate the effects of reactive astrocytes on NSC activation. The control astrocytes (Control As; unstimulated) and reactive astrocytes (Reactive As) or their conditioned medium were co-cultured with NSCs, then the protein aggregates in NSCs and their proliferation capacity were assessed (Figure 3C). After co-culture for 24 hours, reactive astrocytes or their conditioned medium (Reactive ACM), but not the control astrocytes or their ACM (Control ACM), resulted in a mass deposition of aggresome in NSCs (Figure 3, D and E and Supplementary Figure 4, F and G). Moreover, the proliferating capacity of NSCs treated with reactive astrocytes or reactive ACM was significantly diminished, whereas those treated with control astrocytes or their ACM was not affected (Figure 3, F and G and Supplementary Figure 4, H and I). Importantly, reactive astrocytes and their ACM also suppressed the proteasome activity of the cocultured NSCs (Figure 3, H and I), and the protein levels of proteasome 20S and UCHL1 in NSCs were also decreased concomitantly (Figure 3, K-L). Together, these results suggested that reactive astrocytes suppress the activation of NSCs, possibly due to the protein aggresome accumulation resulting from the inhibition of UCHL1-UPS signaling.

**Figure 3.**
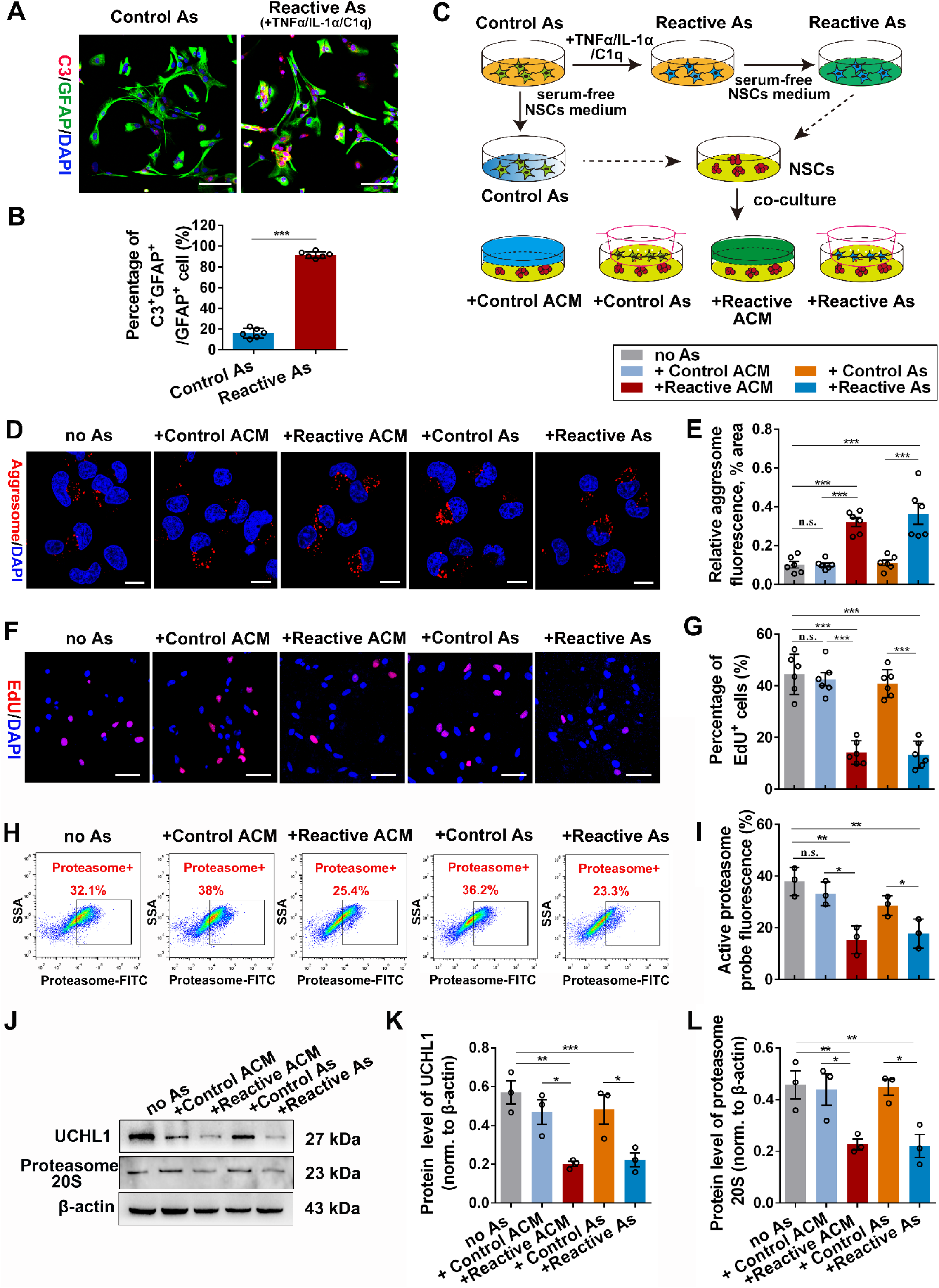
Reactive astrocytes significantly suppressed NSC activation by inhibiting protein aggresome elimination in vitro. (A-B) Immunofluorescent staining showing the successful induction of reactive astrocytes in vitro, which was confirmed by the highly expressed C3^+^ astrocytes. Representative images (A) and quantification (B) of C3^+^ astrocytes were revealed. Scale bar (A), 20 μm. (B) n=6 biological replicates. Data are presented as mean ± SEM. p-values (***p<0.001) are calculated using two-tailed unpaired Student’s t-test. (C) Schematic diagram of the co-culture system of NSCs and astrocytes. (D-E) Confocal representative images (D) and quantification (E) of protein aggregations identified by aggresome dye in NSCs. Scale bar (D), 10 μm. (E) n=6 biological replicates. (F) The activation of NSCs was assessed using EdU incorporation assay. Scale bar (F), 25 μm. (G) The histogram shows the quantification of EdU^+^ NSCs in the co-culture system. n=6 biological replicates. (H) Flow cytometry assay revealed the proteasome activity in NSCs co-cultured with astrocytes for 24h. (I) Quantification of proteasome activity in NSCs co-cultured with astrocytes for 24h. n=3 biological replicates. (J-L) Western blotting analysis of NSCs after co-cultured with astrocytes or ACM. Representative immunoblots and quantification of UCHL1 and 20S proteasome was revealed in J and K/L. (K/L) n=3 biological replicates. (E/G/I/K/L) Data are presented as mean ± SEM. p-values (*p<0.05, **p<0.01, ***p<0.001) are calculated using one-way ANOVA with Tukey HSD post hoc test. See also supplementary Figure 4.

### Reactive astrocytes regulated protein aggregation-associated NSC activation by C3/ C3aR pathway

As part of the innate immune system, the complement system in CNS is emerging to be the important factor mediating the crosstalk between different cell types during development or after injury. For instance, complement factor C3 is obviously activated in the AD brain and involved in neurodegeneration processes (Hong *et al*., 2016; Wu *et al*., 2019). Given that C3 is one of the most prominent class of proteins upregulated in the neurotoxic reactive astrocytes medium (Guttenplan *et al*., 2021), we therefore asked whether astrocytic C3 release is involved in the processes of excessive protein aggregations and impaired activation of NSCs. Consistent with previous studies (Guttenplan *et al*., 2021), increased level of C3a was detected in the medium of NSCs treated with the reactive ACM or reactive astrocytes by ELISA assay (Supplementary Figure 5A). To further investigate the effects of C3 on protein aggregation and activation of NSCs, NSCs were treated with different doses of C3a (0.5, 1 µg/ml). Results revealed that C3a treatment led to increased accumulation of protein aggregations (Figure 4, A and B; Supplementary Figure 5, B and C) and decreased proliferation in NSCs (Figure 4, C and D and Supplementary Figure 5, D and E). In support, the proteasome activity in NSCs was also suppressed by C3a (Figure 4, E and F), which was further confirmed by the decreased expression of UCHL1 (Figure 4, G and H).

**Figure 4.**
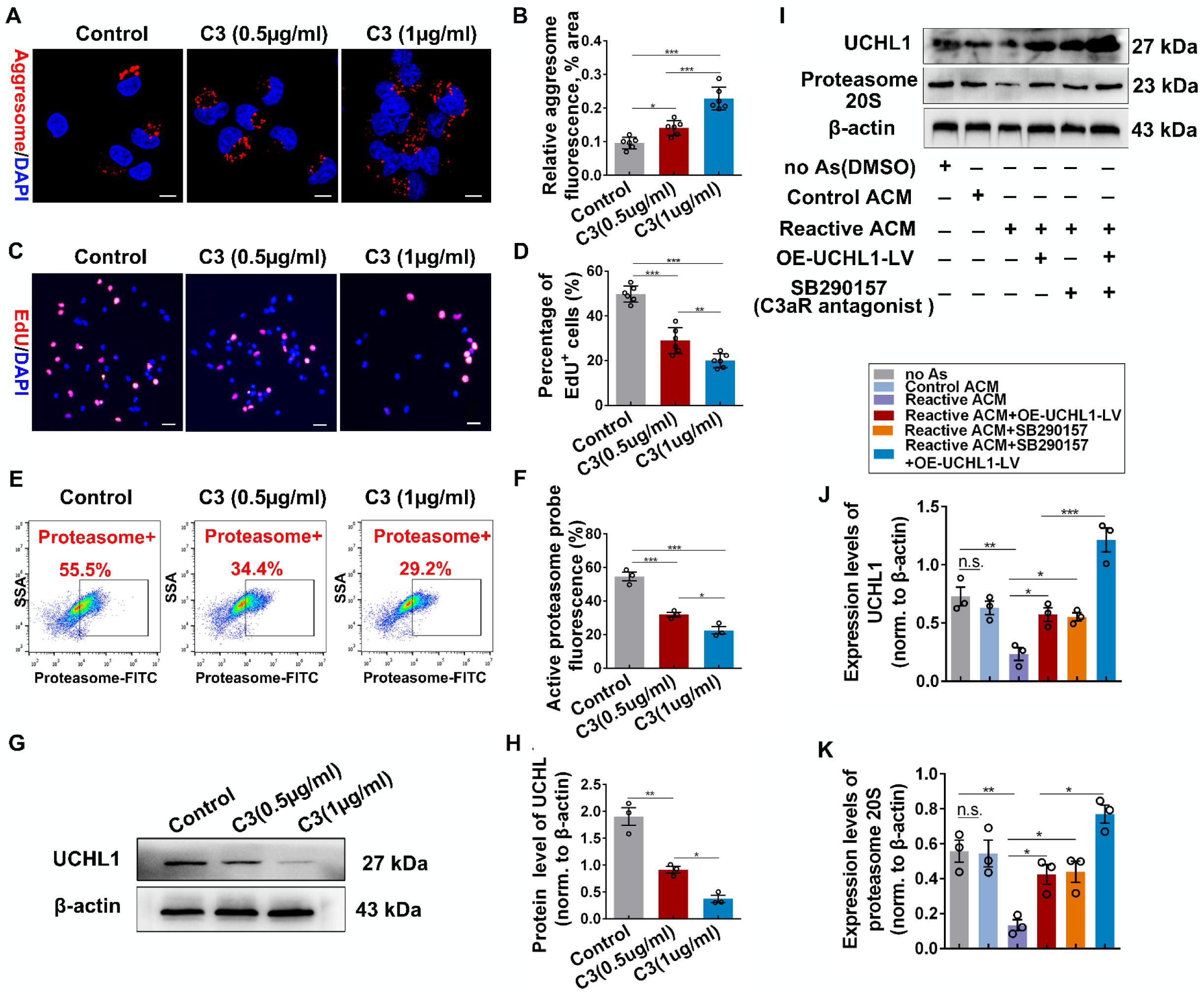
C3 restricted NSC activation by inhibiting UCHL1 and protein aggregation clearance via C3/C3aR pathway. (A-B) Immunofluorescence staining (A) and quantification (B) of protein aggregates (aggresome^+^) in NSCs treated with C3 for 24h. Scale bar (A), 10 μm. (B) n=6 biological replicates. (C) Fluorescent staining with EdU^+^ showed the proliferation of NSCs after cultured with C3 for 24h. Scale bar (C), 25 μm. (D) The percentage of EdU+ NSCs was quantified in different groups. n=6 biological replicates. (E) Flow cytometry assay revealed the proteasome activity in NSCs treated with C3 for 24h. (F) Quantification of proteasome activity by flow cytometry assay in treatments. n=3 biological replicates. (G-H) Expression of UCHL1 and mono-ubiquitin in NSCs incubated with C3 for 24h was measured by Western blotting. Representative images and quantification of proteins were shown in G and H. (H) n=3 biological replicates. (I-K) Western blot analysis of NSCs treated with OE-UCHL1-LV and C3aR antagonist in the reactive astrocytes conditioned medium was conducted with indicated antibodies. Representative immunoblot images and quantification of relative protein level was illustrated in I and J/K. (J/K) n=3 biological replicates. (B/D/F/H/J/K) Data are presented as mean ± SEM. p-values (*p<0.05, **p<0.01, ***p<0.001, n.s. no significant) are calculated using one-way ANOVA with Tukey HSD post hoc test. Source data are provided as a Source data file. See also supplementary Figure 5.

We next examined if complement receptor was required to translate astrocytic C3 release into NSC responses. First, C3a receptor (C3aR) was detected in the cytomembrane of NSCs, similar to that in BV2 microglia expressing C3aR (Supplementary Figure 5, F and G). Second, to explore the connection among reactive astrocytes, C3/C3aR and NSC responses, NSCs were treated with OE-UCHL1-LV or C3aR antagonist (SB290157) in the presence of reactive ACM. Results showed that the decreased levels of UCHL1 and proteasome 20S in NSCs cultured with reactive ACM were all rescued by overexpression of UCHL1 or C3aR blockade (Figure 4, I-K). These evidences support an important role of C3/C3aR signaling in mediating the effects of reactive astrocytes on NSC activation by inhibiting UCHL1 and proteasome activity to impede protein aggregates degradation.

### Up-regulation of UCHL1 enabled endogenous NSC activation in rat spinal cord after SCI

NSCs often remain quiescent in the normal CNS, but they have potential to proliferate and differentiate under trauma condition. However, the activation of spinal cord NSCs in response to SCI is very limited, and neurons are rarely produced in the inhibitory injury microenvironment. To determine the potential effects of UCHL1 on spinal cord Nestin^+^ NSC activation in vivo, the T10 complete transection rat SCI model was performed. The electrophysiological assay was conducted to confirm the successful construction of the complete transection SCI model (Supplementary Figure 6A). NSCs were activated quickly at five days after injury (Supplementary Figure 6, B and C); moreover, many of Nestin^+^ cells also expressed the another NSC marker SOX2 and exhibited shape transformation with hypertrophy and thickening process (Supplementary Figure 6, D and E), suggesting that most of the Nestin^+^ cells might be endogenous NSCs and could be activated in response to SCI.

To manipulate UCHL1 level in the spinal cord, lentiviral vector encoding UCHL1 (OE-UCHL1-LV), recombinant human UCHL1 protein (rh-UCHL1), or the specific UCHL1 inhibitor LDN-57444 was injected into the injury site (Figure 5A). At seven days post-SCI, the remarkably increased level of UCHL1 and proteasome 20S were measured within the damaged spinal cord in OE-UCHL1-LV or rh-UCHL1 group (Figure 5, B-D). We next examined the activation of Nestin^+^ NSCs within the lesion site. More abundant Nestin^+^ cells were found in OE-UCHL1-LV or rh-UCHL1 group at the lesion center than the lesion control, and much fewer Nestin^+^ cells were observed by LDN-57444 treatment (Figure 5, E and F). Moreover, OE-UCHL1-LV or rh-UCHL1 treatment led to obviously increased Ki-67/Nestin^+^ cells, whereas LDN-57444 inhibited NSC proliferation (Figure 5, G and H), indicating that UCHL1 could promote spinal NSC activation after SCI.

**Figure 5.**
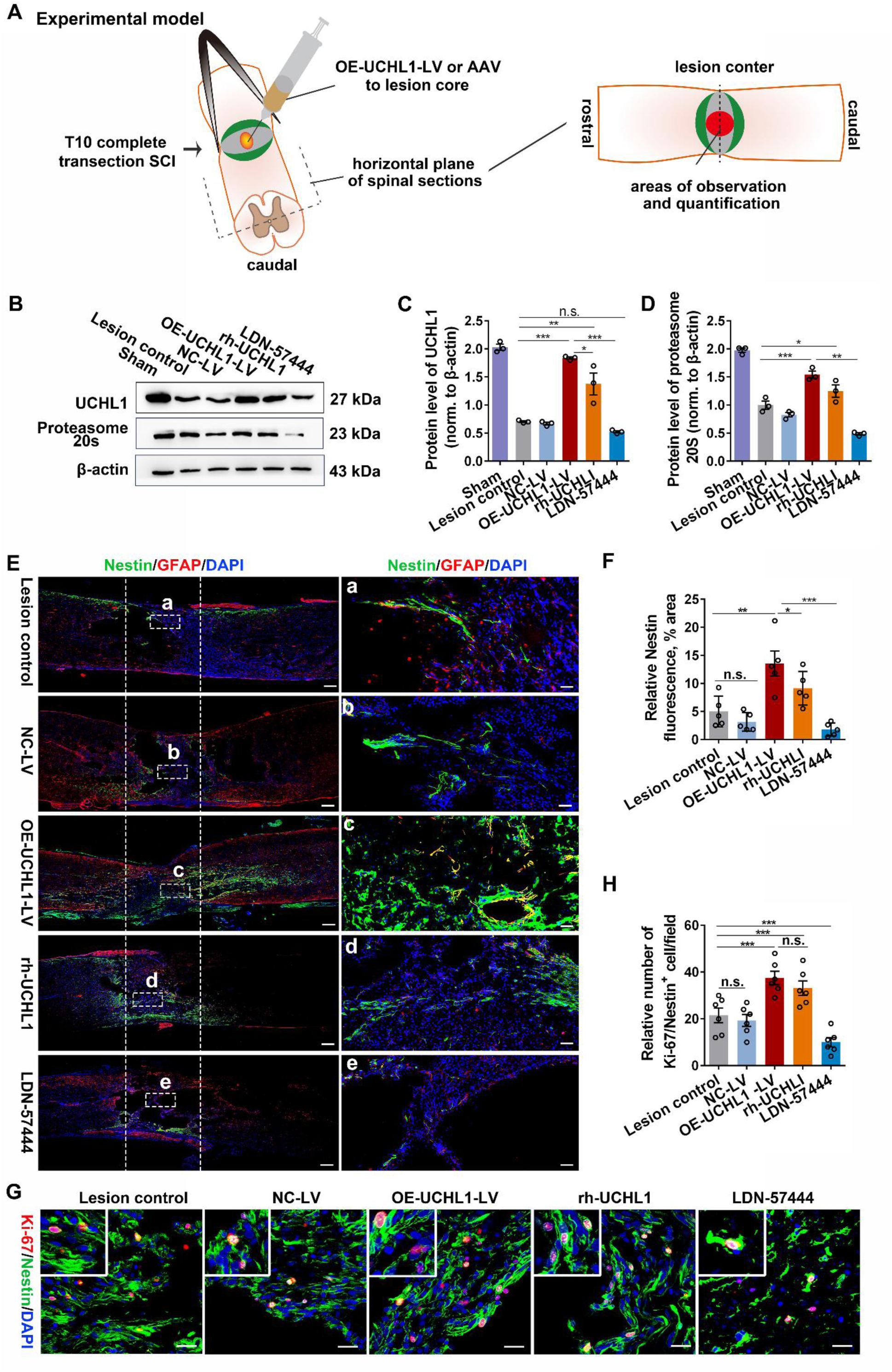
Up-regulation of UCHL1 facilitated the activation of spinal cord NSC after SCI in rats. (A) The diagram illustrates the construction of complete transection SCI model, the injection of the lentivirals or AAV after SCI, and where micrographs are imaged from and the quantitative sites in immunohistochemistry assay. The images for quantification were all captured from the lesion site in SCI animals. (B) The expression of UCHL1 and proteasome 20S in T10 spinal tissues at 7 days after administration of OE-UCHL1-LV, rh-UCHL1 or LDN-57444 in SCI rats by Western blot analysis. (C-D) Quantification of protein levels of UCHL1 and proteasome 20S in spinal tissues at 7 days post-SCI. n=3 independent animals. (E) Confocal representative images showing the expression of Nestin^+^ NSCs and GFAP^+^ astrocytes in spinal cord at 8 weeks after SCI among different treatments. The enlarged images of the white boxed region were illustrated in the right column. The white dotted areas represented the lesion epicenter. Scale bar (C), 50 μm. Scale bar (a-e), 25 μm. (F) Quantification of relative Nestin expression in the lesion center. n=5 independent animals. (G) Confocal representative images show the proliferation of NSCs labeled by Ki-67/Nestin in the lesion core at 7 days after SCI. Scale bar, 20 μm. (H) Quantification of Ki-67/Nestin^+^ cells in Figure A. n=6 independent animals. (C/D/F/H) Data are presented as mean ± SEM. p-values (*p<0.05, **p<0.01, ***p<0.001, n.s. not significant) are calculated using one-way ANOVA with Tukey HSD post hoc test. See also supplementary Figure 6.

To further explore the activation and fate differentiation of Nestin^+^ NSCs in vivo more specifically, we constructed the tdTomato tagged adeno-associated virus (AAV) containing the Nestin-specific promoter to achieve the targeted overexpression of UCHL1 in Nestin^+^ NSCs (Figure 6A). The expression of UCHL1 within the damaged spinal segments was detected at two weeks later after administration of OE-UCHL1-AAV in SCI rats (Supplementary Figure 6, F and G). Importantly, about 82.57% Nestin^+^ cells were co-labeled with tdTomato in either NC-AAV or OE-UCHL1-AAV group (Supplementary Figure 6H-I), indicating the high infection efficiency and specificity of AAV targeted in Nestin^+^ cells in vivo. To evaluate the Nestin^+^ NSC activation after SCI, rats were intraperitoneally injected with BrdU daily post-injury for two weeks. Compared with the control group, OE-UCHL1-AAV significantly promoted Nestin^+^ NSC activation (Figure 6, B and C), in consistent with the previous results obtained with the Lenti-viral vectors (Figure 5, E-H). Moreover, some tdTomato/Tubulin β3^+^ cells and tdTomato/DCX^+^ cells were detected at the lesion site of OE-UCHL1-AAV infected rats (Figure 6, D-G), suggesting the potential possibilities of the newborn neuron generation and neurogenesis from the manipulated Nestin^+^ NSCs. More importantly, a few BrdU/Tubulin β3^+^ and BrdU/NeuN^+^ neurons were observed in the OE-UCHL1-AAV group (Figure 6, H-J), implying certain generation of newly-born neurons from the proliferating Nestin^+^ NSCs. Nevertheless, what kind of neurons they may be formed, or where those neurons may project and how they may contribute to functions is still unknown and worthy of further study.

**Figure 6.**
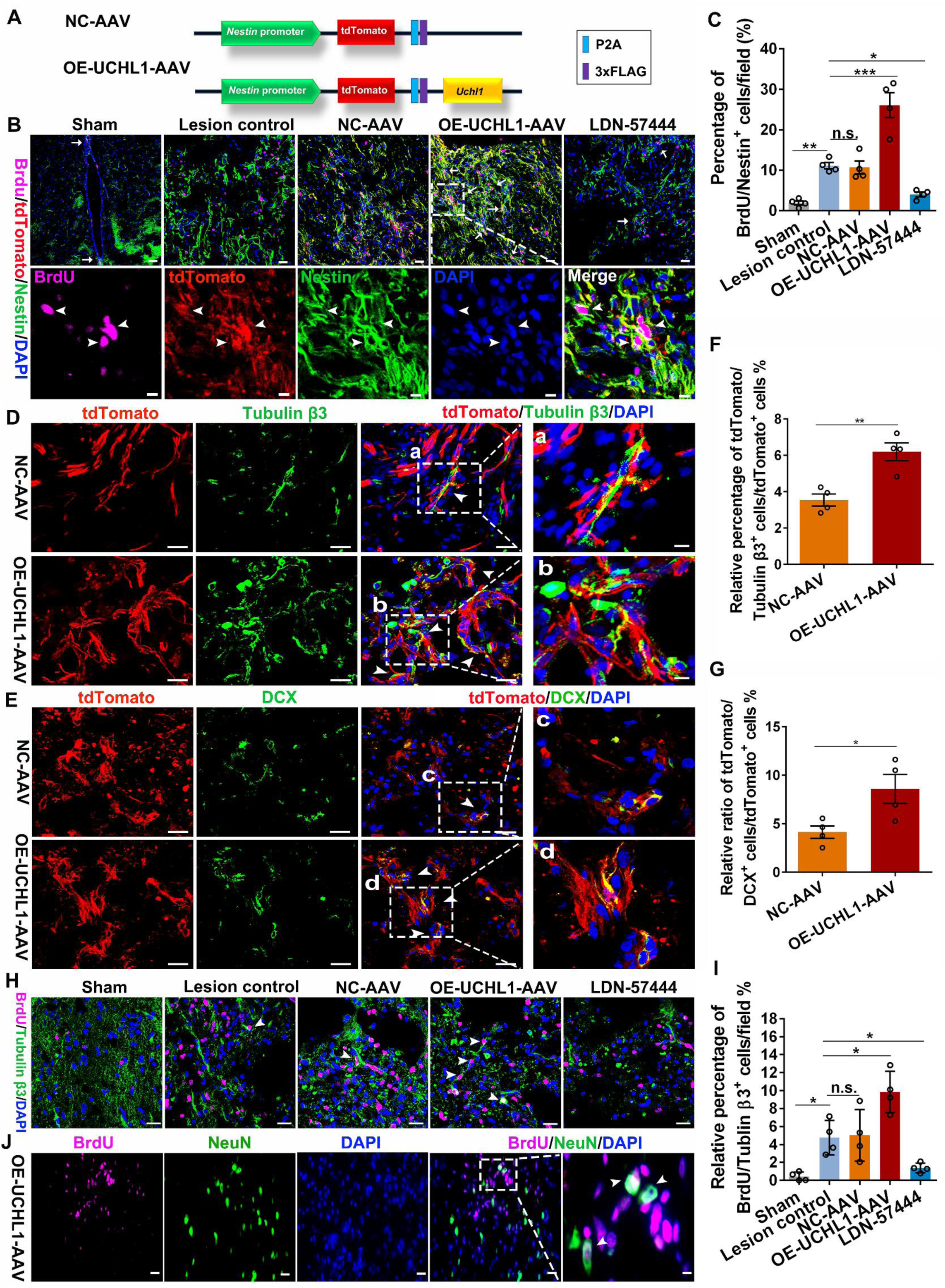
Overexpression of UCHL1 using OE-UCHL1-tdTomato AAV promoted NSC proliferation post-SCI. (A) Schematic model of AAV containing the specific Nestin promoter to target overexpression of UCHL1. (B) Confocal representative images show the proliferation of NSCs labeled by BrdU and Nestin in the lesion center at two weeks after treatment of OE-UCHL1-AAV in SCI rats. The white arrow indicated BrdU/Nestin^+^ cells. Scale bar (the up row), 20 μm. Scale bar (the down row), 10 μm. (C) Quantification of BrdU/Nestin^+^ NSCs in the lesion at two weeks after SCI. n=3 independent animals. (D-E) TdTomato positive neurons around the lesion were possibly differentiated from the activated endogenous NSCs. The white triangle indicated tdTomato/Tubulin β3^+^ cells and tdTomato/DCX^+^ cells. Scare bar, 20 µm. Scare bar (a-d), 10 µm. (F-G) Quantification of tdTomato/Tubulin β3^+^ cells (F) and tdTomato/DCX^+^ cells (G) in the lesion at two weeks after SCI. n=3 independent animals. (H) Newborn neurons within the lesion center were identified by immunofluorescence staining of BrdU/Tubulin β3^+^ at two weeks post-SCI. The white triangle indicated BrdU/Tubulin β3^+^ cells. Scale bar, 20μm. (I) Quantification of BrdU/Tubulin β3^+^ neurons in the lesion at two weeks after SCI. n=3 independent animals. (J) Some BrdU/NeuN^+^ neurons were observed in the lesion of OE-UCHL1-AAV rats. Scale bar (H), 20 μm. Scale bar (a), 10 μm. Data are presented as mean ± SEM. p-values (*p<0.05, **p<0.01, ***p<0.001, n.s. not significant) are calculated using (C/I) one-way ANOVA with Tukey HSD post hoc test or (F/G) two-tailed unpaired Student’s t-test.

In addition, improved locomotor functions were detected in SCI rats treated with OE-UCHL1-LV or rh-UCHL1 by BBB score test (Basso et al., 1995) from the 5th to 8th weeks post-SCI, and blocking UCHL1 activity with LDN-57444 impaired the spontaneous recovery of hind limbs (Supplementary Figure 6J). The electrophysiological assay was further performed to evaluate the locomotor functional restoration before perfusion. A stimulating electrode was positioned at the intraspinal dorsal T7 spinal cord (3 segments above the lesion), and any evoked activity at T13 spinal cord (3 segments below the lesion) was recorded (Supplementary Figure 6K). In the uninjured animals, a short latency response was evoked by stimulation at T7, which was entirely abolished after T10 transection (Supplementary Figure 6A). After 8 weeks, evoked response was partially restored in SCI animals treated with OE-UCHL1-LV or rh-UCHL1, implying certain reconnection of synaptic relays in the lesion epicenter (Supplementary Figure 6L and M). It is speculated that the activated NSCs may underlie the neurological repair by bridging the disconnected spinal cord after SCI. However, we cannot rule out the possibility that UCHL1 could also have positive effects on residential spinal neurons and their axons, which also could contribute to the observed motor function recovery. Therefore, the potential mechanisms contributing to the functional improvement is not very clear and needs further investigation.

### Blockade of reactive astrocytes and C3/C3aR pathway promoted NSC activation in SCI mice

The above studies demonstrated the important roles of neurotoxic reactive astrocytes in NSC activation. We next assessed whether inhibition of reactive astrocytes affected endogenous NSC activation after SCI in vivo. Due to the high dosage of neutralizing antibodies required in vivo, C57BL/6 mice rather than SD rats were selected to conduct T10 transection experiment. SCI mice were administrated with neutralizing antibodies against IL-1α, TNFα and C1q to inhibit reactive astrocytes formation as previously reported (Liddelow *et al*., 2017). Massive reactive astrocytes were induced at seven days post-SCI, and neutralizing antibodies, but not the IgG isotype control, successfully blocked reactive astrocyte formation (Supplementary Figure 7, A and B). Enhanced proliferation of NSCs (Ki67/Nestin^+^) around the central canal and in the lesion epicenter were induced by neutralizing antibody administration (Supplementary Figure 7, C and D). Western blot analysis revealed that blockade of reactive astrocytes led to decreased level of C3a, and increased expression of UCHL1 and proteasome 20S (Supplementary Figure 7, E-H). No significant difference was detected between the lesion control and the IgG treated groups. To further investigate if reactive astrocytes mediated NSC activation by C3/C3aR pathway, Sham and SCI mice were administrated with C3aR antagonist (SB290157) or 0.9% saline intraperitoneally daily. At seven days post-injury, compared with the lesion control group, the EdU-positive Nestin^+^ NSCs were significantly increased when C3/C3aR signaling was blocked (Figure 7, A and B), as well as the expression of Nestin, UCHL1 and proteasome 20S (Figure 7, C and F). Our data showed that blockade of reactive astrocytes might facilitate Nestin^+^ NSC activation post-SCI, possibly via C3/C3a/UCHL1 pathway. Nevertheless, due to these three cytokines and C3aR are also expressed on other cells in addition to astrocytes, the neutralizing antibodies or C3aR antagonist may also affect NSC activation through other potential approaches such as alleviating the neuroinflammatory microenvironment.

**Figure 7.**
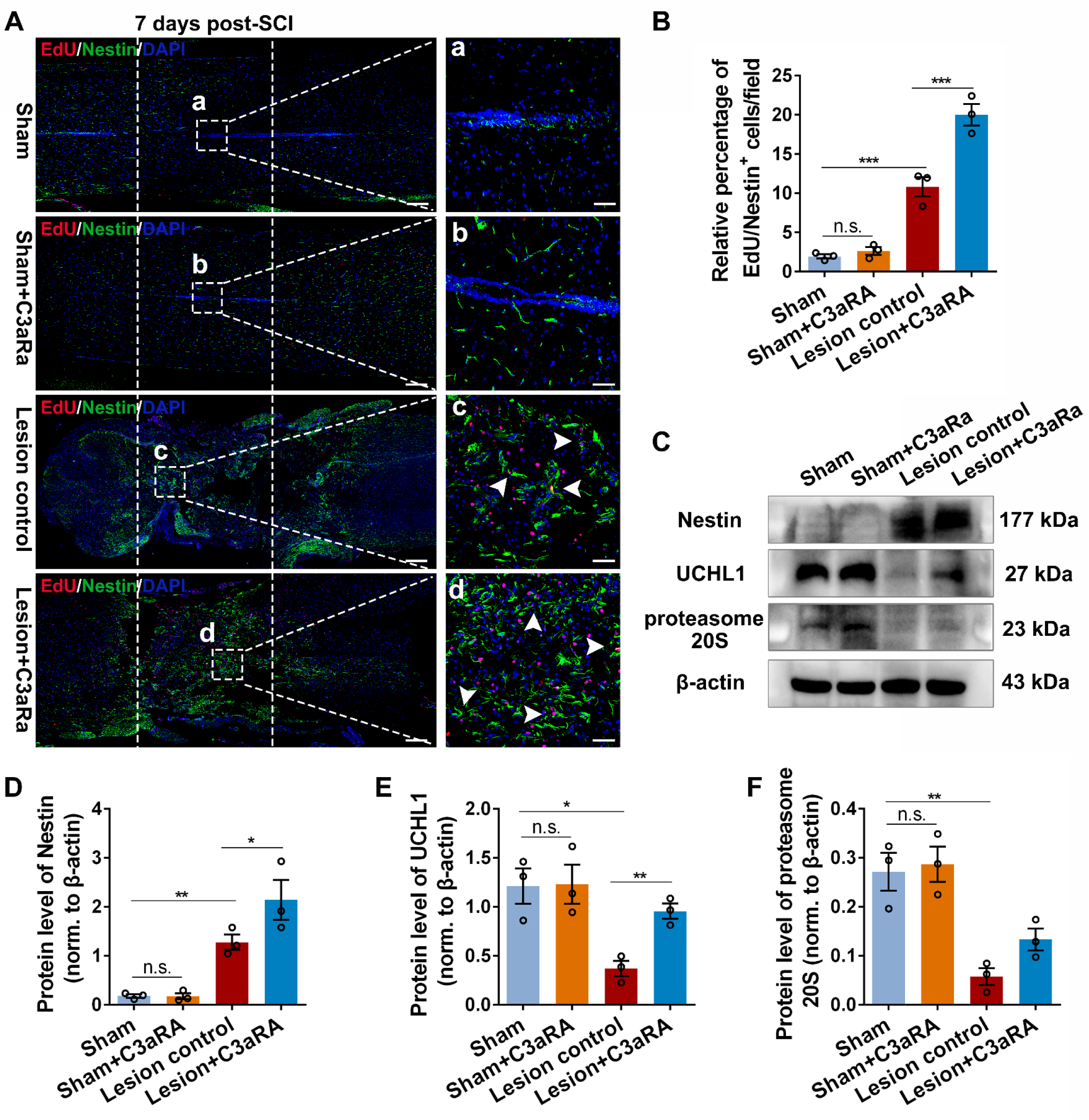
Blockade of C3/C3aR pathway enhanced NSC activation in lesion site after SCI. (A) Confocal representative images show the proliferation of NSCs labeled by EdU/Nestin in the lesion core at 7 days after SCI. Scale bar, 20 μm. (B) Quantification of EdU/Nestin^+^ cells in Figure A. n=3 independent animals. (C) The expression of Nestin, UCHL1 and proteasome 20S in T10 spinal tissues at 7 days after administration of C3aR antagonist in SCI mice by Western blot analysis. (D-F) Quantification of protein levels of Nestin, UCHL1 and proteasome 20S in spinal tissues at 7 days post-SCI. n=3 independent animals. (B/D/E/F) Data are presented as mean ± SEM. p-values (*p<0.05, **p<0.01, ***p<0.001, n.s. not significant) are calculated using one-way ANOVA with Tukey HSD post hoc test.

## DISCUSSION

Neuronal cell death is the major reason underlying the loss of sensory, motor, and cognitive functions after neural injuries or neurodegenerative diseases. Replacement of lost neurons can be achieved through cell transplantation(Lu et al., 2012; Salewski et al., 2015), in vivo direct reprogramming for glia-neuron trans-differentiation(Li and Chen, 2016; Puls et al., 2020; Tai et al., 2021), or activation of endogenous NSCs(Goncalves et al., 2016; Maeda et al., 2019; Stenudd et al., 2015). However, emerging stringent lineage tracing study(Wang et al., 2021) has recently challenged and raised the bar of evidence for direct reprogramming astrocytes into neurons in vivo. Limited number of NSCs still remain in the mature CNS. Although activation and neuronal differentiation of endogenous NSCs have been reported in the brain hippocampus (Denoth-Lippuner and Jessberger, 2021; Goncalves *et al*., 2016), the NSCs in the adult spinal cord often fail to do so. One likely reason for such failure in the spinal cord is that the local microenvironment does not support the endogenous NSC activation and neuronal fate.

In this study, we identified UCHL1 as a key positive regulator of spinal cord Nestin^+^ NSC activation. Specifically, SCI led to down-regulation of UCHL1 in NSCs through C3 produced by neurotoxic reactive astrocytes. Therefore, direct overexpression of UCHL1 in the NSCs resulted in their activation via the enhanced ubiquitin-proteasome-mediated clearing of intracellular protein aggregates. As a result, overexpression of UCHL1 or blockade of reactive astrocytes and C3/C3aR pathway enhanced endogenous Nestin^+^ NSC activation after SCI. Our findings revealed that the Nestin^+^ spinal cord NSCs were negatively controlled by reactive astrocyte-mediated inactivation of the UCHL1-proteasome pathway, therefore providing new insights into the non-neurogenic pathological milieu of mature spinal cord after SCI.

It has remained highly contentious concerning the resident NSCs in adult spinal cord. Ependymal cells are suggested to be the latent NSCs with neurogenic functions(Llorens-Bobadilla et al., 2020; Meletis *et al*., 2008), which is now disputable in view of the contradictory results from fate mapping and single-cell spatial transcriptomics analysis (Shah *et al*., 2018; Shu *et al*., 2021; Stenudd et al., 2022). Nestin^+^ cells in the spinal cord proliferate quickly post-SCI and most of them also show diversified gene expression patten similar with several types of cells such as neurons, astrocytes and oligodendrocytes in addition to ependymal cell(Meletis *et al*., 2008; Shu *et al*., 2021), suggesting their stem cell-like properties of self-renewal and multipotency after injury. Nestin/SOX2^+^ cells are commonly identified as the neural stem/progenitor cells in CNS (Chen et al., 2022; Defterali et al., 2021; Morrow *et al*., 2020). In the complete transection SCI model here, majority of Nestin^+^ cells activated after injury were also colocalized with the NSC marker SOX2, which we thus consider as the endogenous NSCs in this study.

Insoluble protein aggregations are important factors involved in regulation of stem cell activation, proliferation and differentiation. Protein homeostasis of NSCs and progenitor cells are proteolytically processed by UPS, in which several ubiquitin ligases and deubiquitinating enzymes, including UCHL1, play essential roles (Gong et al., 2016). A previous study showed that quiescent NSCs could utilize vimentin-mediated proteasome localization to clear protein aggresome during activation, and Nestin^+^ NSCs in vimentin KO mice have decreased ability to exit quiescence status in vivo (Morrow *et al*., 2020). It has also been reported that the removal of protein aggregates by enhancing lysosome activity improves the capacity of quiescent NSCs to be activated (Leeman *et al*., 2018). In the present study, overexpression of UCHL1 in NSCs facilitated their proliferation similarly through the proteasome-mediated protein aggregations clearance in vitro. Importantly, targeted upregulation of UCHL1 also enhanced the activation of Nestin^+^ NSCs response to injury in adult SCI animals. Moreover, discoveries (Sakurai *et al*., 2006) suggested that UCHL1 promote neuronal differentiation and neurogenesis of Nestin^+^ progenitors in the embryonic brain. Interestingly, a few newborn neurons (BrdU/Tubulin β3^+^ and BrdU/NeuN^+^ neurons) were detected in the injured spinal cord here after UCHL1 upregulation in Nestin^+^ NSCs by AAV. However, whether these cells could form functional neurons, or whether and how they may contribute to the nerve circuit reconstruction are unclear for further exploration. NSCs derived from the brain or adult spinal cord exhibit similar properties of self-renewing and multipotency in vitro. In view of the difficulties in cell extraction, purification, and limited number of NSCs in adult spinal cord, fetal NSCs from the brain, which were widely used in plentiful stem cell researches, were utilized here in vitro experiments. Although certain differences between the fetal and adult NSCs, our data show that UCHL1 enhanced both of their activation in vitro and in vivo separately, which support the pivotal role of UCHL1-related UPP in NSCs fate mediation. The potential mechanism underlying the promotive role of UCHL1 on NSCs may largely attribute to its regulatory effects on ubiquitin-proteasome activity as previously described (Naujokat, 2009b; Sakurai *et al*., 2006).

In addition, UCHL1 plays critical role on axonal integrity, axonal transport and synaptic functions. It protects neurons from death by removing damaged proteins via UPP, preserves axon functions and improves sensorimotor recovery after CNS injury (Gong *et al*., 2006; Liu *et al*., 2019). Here, the improved locomotor functions were also observed in SCI rats after UCHL1 upregulation. Several factors might underlie the UCHL1-induced improvement of neural repair after SCI. One potential factor is that newly generated neurons from NSCs at the injury site might partially contribute to the neural reconnection by extending axons towards both directions of the spinal cord. Nevertheless, whether and how much of the detected functional improvement result from the activated NSCs and their subsequent effects are difficult to ascertain. Furthermore, the potential protection effects of UCHL1 on local spinal neurons, the regeneration of injured descending axons, and/or the sprouting from spared uninjured axons are all likely involved in the improved functional outcomes. Therefore, further investigations are required to clarify the potential mechanisms by functional recovery is achieved.

The molecular mechanism underlying impaired proteasome and UCHL1 function in the spinal cord NSCs is mostly unknown. Based on protein microarray analysis of CSF, we found that C3 and complement cascades pathway were remarkably upregulated in SCI animals. Moreover, C3^+^ reactive astrocytes were abundantly activated in the lesion site after SCI. Reactive astrocytes are neurotoxic and can propagate inflammatory signals during neurodegeneration (Guttenplan *et al*., 2021; Liddelow *et al*., 2017). Their inappropriate activation upregulates the classical C1q/C3 complement pathway, resulting in the impaired clearance of cellular debris and the formation of amyloid β (Aβ) plaques in aging or AD related neurons (Alawieh et al., 2015; Hong *et al*., 2016; Rogers et al., 2002; Yun et al., 2018). In this study, reactive astrocytes induced by TNFα/IL-1α/C1q significantly suppressed NSC proliferation by impeding protein aggregations clearance. Evidence from the administration of C3aR antagonist to NSCs cultured with reactive ACM revealed that C3/C3aR signaling likely mediated the suppression of UCHL1 and UPS activity by reactive astrocytes, eventually resulting in protein aggregation-associated dysfunction of NSC activation. Furthermore, abolishment of reactive astrocytes with neutralizing antibodies or the blockade of C3/C3aR pathway could facilitate NSC activation after SCI, possibly through UCHL1-proteasome pathway. Considering the complex microenvironment in vivo and the nonspecific-expression of C3aR in other cells, it cannot be overlooked that the neutralizing antibodies and C3/C3aR signaling blocking may also regulated the NSC activation through other approaches, not only by the mechanism of UCHL1-UPP-mediated aggregates clearance.

Activation of endogenous NSCs is a challenging task of great importance to enhance intrinsic neural regeneration after CNS injury. Uncovering of the mechanisms underlying the complicated cellular and molecular interactions within NSCs microenvironment is the critical basics for the therapeutic strategies. Our study suggested that reactive astrocytes activated by SCI might be one of the local environmental factors contributing to the non-neurogenic environment in the adult spinal cord, which inhibits UCHL1 and impairs UPS functions in NSCs to impede their activation through the C3/C3aR signaling. The present study revealed that neurotoxic reactive astrocytes, C3 signaling, and UCHL1 associated proteasome system could all be potential therapeutic targets for NSC activation after SCI.

### Limitations of the study

The present study mainly focused on the regulatory mechanism underlying the activation of spinal cord NSCs post-SCI. Although it remains debating what cells are considered endogenous NSCs in the spinal cord, here we identified the endogenous NSCs as Nestin^+^ cells, majority of which are also SOX2^+^. To more specifically target and trace fate determination of endogenous NSCs for genetic manipulation in the adult spinal cord, fate mapping or single-cell spatial transcriptomics analysis are needed to identify which type of cells serving as the endogenous NSCs. Additionally, gene knockout technology is necessary to more clearly elucidate the regulation of reactive astrocytes on NSC activation in the adult spinal cord.

## ACKNOWLEDGMENTS

The study was supported (to D.Y.B.D.) by National Natural Science Foundation of China (NSFC; 82071362), Basic Research Project of Shenzhen Science and Technology Innovation Commission (JCYJ20190809165201646; JCYJ20210324123001003; JCYJ20220530144801003), and Scientific research project of Traditional Chinese Medicine in Guangdong Province (20221096). We thank Krsna Muscheck and Changlin Zhang for helpful suggestions and revision on the manuscript.

## AUTHOR CONTRIBUTIONS

L.D., W.W.C. and D.Y.B.D. designed the experiments. L.D., W.W.C. performed the experiments and analyzed the data. X.Y., and M.S. helped with the behavior tests and histology experiments. L.D. wrote the manuscript. W.W.C., T.L., F.-Q.Z. and D.Y.B.D. revised the paper. All authors reviewed and approved the manuscript.

## STAR★METHODS

Detailed methods are provided in the online version of this paper and include the following:

- KEY RESOURCES TABLE
- RESOURCE AVAILABILITY

- Lead Contact
- Materials Availability
- Data and Code Availability
- EXPERIMENTAL MODEL AND SUBJECT DETAILS

- animals
- METHOD DETAILS

- NSCs Culture and Differentiation
- Astrocytes Culture and Induction
- Lentiviral-mediated overexpression of UCHL1
- Co-culture of NSCs and Astrocytes
- Cell Proliferation Assay
- Proteostat Analysis
- Proteasome Activity Assay
- Enzyme Linked Immunosorbent Assay (ELISA)
- Spinal Cord Injury Model and Lentivirus Administration
- Adeno-associated Virus (AAV) Construction and Injection
- BrdU Assay
- Blockade of Reactive Astrocytes and C3/C3aR pathway in SCI Mice
- Behavior Test
- Electrophysiology Examination
- Protein Microarray of Cerebrospinal Fluid from SCI Animals
- Tissue Preparation
- Immunofluorescent Staining
- Quantitative Real-time Polymerase Chain Reaction (qPCR)
- Western Blotting
- QUANTIFICATION AND STATISTICAL ANALYSIS

- Quantification of Confocal Analysis
- Statistics

### KEY RESOURCES TABLE

### EXPERIMENTAL MODEL AND SUBJECT DETAILS

#### Animals

Female Sprague Dawley (SD) rats (age, 8∼10 weeks; weight, 180∼200 g) and C57BL/6 mice (half male and female; age, 8∼10 weeks; weight, 15∼20 g) were purchased from the Guangdong Medical Laboratory Animal Center and housed under standard specific pathogen free (SPF) condition. All animals were housed in standard cages under a regulated environment (12-h light/dark cycle) with free access to food and water about one week before surgery. All animal experimental procedures using laboratory animals were conducted in accordance with the Guide for the Care and Use of Laboratory Animals (National Research Council, 1996) and approved by the Animal Care and Use Committee of Sun Yat-sen University (SYSU-IACUC-2021-000438).

### METHOD DETAILS

#### NSCs Culture and Differentiation

NSCs were obtained from the fetal brain of embryonic14-day SD rats. Briefly, the cerebrum of embryos was dissected out and the covering pia mater and blood vessels were removed under a microscope. Then the brain was mechanically dissociated in pre-cooling phosphate-buffered saline (PBS) to generate a single cell suspension and centrifuged at 1000 revolutions per minute (rpm) for 10 min. The cell pellet was resuspended and cultured with DMEM/F-12 medium (Gibco, Life Technologies, USA) supplemented with 2% B-27™ supplement (50X; Gibco, Life Technologies, USA), 20 ng/ml growth factor (EGF; Peprotech, New Jersey, USA), 20 ng/ml fibroblast growth factor (FGF; Peprotech, New Jersey, USA), and 1% penicillin/streptomycin (10,000 U/ml, Gibco, Life Technologies, USA). NSCs were purified by passaged every three days, and cells between passages 2∼5 were selected to perform further investigation.

For NSCs differentiation, the cells pellets were digested into single cells with Accutase (Millipore, Bedford, MA), and subjected to neuronal differentiation on 0.1% poly-L-lysine (Gibco, USA) coated culture dishes in Neurobasal (Gibco, Life Technologies, USA) supplemented with 2% B-27™. The medium was changed every 2-3 days.

#### Astrocytes Culture and Induction

Astrocytes were isolated from postnatal two-day SD rats. After stripping the meninges and blood vessels, cortices were mechanically then enzymatically dissociated into single-cell suspension with 0.25% trypsin at 37°C for 15 min, followed by filtration and centrifugation to collect cell pellets. Cells were suspended with basal medium (DMEM/F-12; 10% fetal bovine serum, FBS) and plated on uncoated plates for 30 min to remove fibroblasts and endothelial cells. Subsequently, the non-adherent astrocytes were transferred into new dishes pre-coated with 0.1% poly-L-lysine. The cell medium was changed every three days and passaged when the cell fusion rate reached approximately 90%. Astrocytes were shaken at 200 r/min overnight to remove upper layer microglia that were attached to the surface of astrocytes, followed by administrated with PLX5622, a specific microglia scavenger, to further purify astrocytes before passage. To induce reactive astrocytes, cells were incubated with the triple factors TNFα (30 ng/ml), IL-1α (3 ng/ml) and C1q (400 ng/ml) for 24 h and the successful induction of reactive astrocytes was verified by highly increased C3^+^ astrocytes formation via immunofluorescent staining and flow cytometry assay.

#### Lentivirus-mediated Overexpression of UCHL1

The lentivirus overexpressing UCHL1 (OE-UCHL1-LV) and empty lentiviral vector (NC-LV) were constructed by HanBio Technology (Shanghai, China). Lentivirus designed to overexpress UCHL1 were cloned into the pHBLV-CMV-MCS-3FLAG-EF1-GFP-T2A-PURO lentiviral vector containing the ZsGreen reporter gene. The lentivirus vectors were transferred into 293T cells in the presence of packaging plasmids (psPAX2 and pMD2G) using LipofiterTM (HanBio Technology) for lentivirus packaging. 293T cells were transfected and the overexpression efficacy were assessed after 48 h by qPCR. The final lentiviral vector titer was 2 × 10^8^ TU/ml.

For NSCs infection, cells were transfected with diluted lentiviral solution with a multiplicity of infection (MOI) of 25, with supplement of 1 mg/ml polybrene. At 24 h after transfection, the medium was replaced with fresh basal medium (FBS-free) without penicillin/streptomycin for further 24 h. GFP expression was visualized under fluorescent microscope at 48 h post-transfection, and over-expression of UCHL1 was determined by Western blot and qPCR analysis.

#### Co-culture of NSCs and Astrocytes

Two in vitro co-culture systems were conducted here (Figure 4E): I. We used a transwell system that allowed interaction via diffusible factors. Unstimulated Control astrocytes or Reactive astrocytes in the transwell were on the top of NSCs in the lower chamber, and both cells were cultured within the basal culture medium (free of FBS) of NSCs. II. a simple astrocytes conditional medium (ACM) transfer from the Control/Reactive astrocytes cultures to NSCs’ cultures. After incubation in the normal or A1-inducing medium, astrocytes were cultured with the basal culture medium (free of FBS) of NSCs for another 24h, then the ACM from the Control astrocytes or Reactive astrocytes were collected and added into the basal medium of NSCs at a ratio of 1:1 ACM to NSCs culture medium. The cell proliferation, aggresome formation and proteasome activity was assessed at 24 h later.

#### Cell Proliferation Assay

Cell proliferation in vitro was examined by EdU incorporation. NSCs were incubated with a culture medium supplemented with 10 µM EdU overnight before harvesting. Cells were fixed with 4% paraformaldehyde (PFA) followed by permeabilization, then washed twice with 3% bovine serum albumin (BSA) and stained using Click-iT EdU assay kit (US Everbright INC.) in accordance to the manufacturer’s instructions. After washed once, the cells were resuspended with Hoechst solution (2 ug/ml; US EVERBRIGHT INC.) for DNA counterstaining prior to analysis on Cytoflex LX or laser scanning confocal microscope (LSCM). The control cells that were not subjected to EdU but underwent fluorescent EdU detection were used as a negative base to assess cutoff values for EdU positivity. For quantification of EdU assay, the ratio of the EdU^+^ cells in at least three replicates among different groups were counted by a blinded observer.

#### Proteostat Analysis

Cells were fixed as mentioned above, then permeabilized with 0.5% Triton X-100/PBS on ice and gently shake for 30 min. After washed twice with PBS, cells were resuspended with PROTEOSTAT Aggresome Detection Reagent (1:2000) and protected from light at room temperature (RT) for 30 min, and subsequently counterstained with Hoechst 33342 (1:1000). The stained cells were analyzed using confocal microscope. For flow cytometry, 500 µl of freshly diluted 1:10000 PROTEOSTAT Aggresome Detection Reagent was applied to incubate cells for 30 min under protection from light. A nutrient deprivation assay that cleared proteostat and a proteasome inhibitor were conducted as controls. MG132, a proteasome inhibitor accelerating the formation of perinuclear aggresome within cells, was used as a positive control. For nutrient deprivation, cells were incubated in HBSS (Gibco) supplemented with 1 mM HEPES that prevented over-acidification for 3 h, then the medium was replaced with basal culture medium. Proteostat labeling was determined by the relative fluorescence area of protein aggresome in the same field of view across different treatments with blind evaluation, with at least three biological replicates.

#### Proteasome Activity Assay

Proteasome activity assays were performed by a proteasome activity probe according to the instructions. The treated cells were incubated with the 5 µM proteasome activity probe Me4BodipyFl-Ahx3Leu3VS (Boston Biochem) for 2 h at 37°C, then washed and analyzed via flow cytometry. The experiment was repeated three times and quantified by a blinded observer.

#### Enzyme Linked Immunosorbent Assay (ELISA)

The C3a level in the peripheral blood and CSF collected from SCI rats were detected through RayBio huMan C3a ELISA kit (RayBiotech) following the manufacturer’s instructions. Briefly, 100 µl standard or sample was added to each well and incubated 2.5 h at RT with gentle shaking, then washed four times and incubated with biotinylated antibody for 1h. 100 µl HRP-conjugated Streptavidin was subsequently added, followed by incubation of 3,3,5,5’-tetramethylbenzidine (TMB) one-step substrate reagent for 30 min at RT in the dark with gentle shaking. Finally, terminating the action with stop solution and the absorbance value at 450 nm was measured and recorded immediately.

#### Spinal Cord Injury Model and Lentivirus Administration

To produce rat model of the 10th thoracic vertebra (T10) complete transection SCI, female SD rats were anesthetized and the back fur was shaved and disinfected with Iodine Volts Swab. An approximately 2 cm incision was made on the T9-T11 skin, then the fat, fascia layer and paravertebral muscles were separated successively to expose the T10 vertebra. Then T10 laminectomy was performed and T10 spinal tissue was transected completely using a sharp scalpel. After hemostasis, rats were randomly divided into six groups, and a total of 10 µl specific mixture was injected into the lesion site using a microsyringe according to the animal groups as follows: Sham group (no SCI; n=12); lesion control group (PBS; n=12); empty lentiviral vector (NC-LV; 2×108 TU/ml; n=12); lentiviral vector encoding UCHL1 (OE-UCHL1-LV; 2×108 TU/ml; n=12) group; recombinant human UCHL1 (rh-UCHL1; 4 µM; n=12) group; and the LDN-57444 (UCHL1 inhibitor; 2 mM; n=12) group. A total of 5 µl virus or recombinant protein was separately mixed with 5 µl Matrigel before injected into the lesions of SCI rats to avoid loss.

The body temperature of animals was maintained at about 37°C using a heating pad during the entire surgery until revival fully from anesthesia. During postoperative care, animals underwent manual bladder evacuations till autonomous urination restoration, and checks daily for wounds, infection, weight loss, autophagy of toes and mobility. All animals used in this experiment did not show wound infection, erosion or self-induced wounds.

#### Adeno-associated Virus (AAV) Construction and Injection

The AAV targeting UCHL1 pAAV-Nestin-tdTomato-P2A-3xFLAG-Uchl1-tWPA or the control vector only with tdTomato fluorescence (pAAV-Nestin-tdTomato-P2A-3xFLAG-MCS-tWPA) were generated by OBiO Technology Corp., Ltd (Shanghai). UCHL1 gene was sub-cloned into pAAV-Nestin-tdTomato-P2A-3xFLAG-WPRE plasmids to produce pAAV-Nestin-tdTomato-P2A-3xFLAG-Uchl1-tWPA. The virus titer was 6.81E+12 Vector Genomes per mL (VG/ml) determined by quantitative PCR. The virus (1.5 µl in volume) was delivered into the lesion center with a microsyringe immediately after SCI and experiments were carried out at two weeks after virus injection to allow the gene expression.

#### BrdU Assay

The activation of NSCs in the spinal cord was evaluated by BrdU (5-bromodeoxy-2′-deoxyuridine; Sigma) after AAV injection. SCI rats administrated with AAV were intraperitoneally injected with BrdU (30 mg/kg body weight) daily after surgery for two weeks until euthanasia. The injured spinal cords were collected and frozen sections (10 µm) were prepared on a cryostat. Sections were pretreated with 2M HCL for 30 min at 37°C, followed by 0.1 M boric acid (pH 8.5) (Biosharp) for 10 min at room temperature. Then sections were blocked, incubated with the anti-BrdU and anti-Nestin primary antibody (1:1000; Sigma) over night at 4°C and Alexa Flour 488/647-conjugated secondary antibodies subsequently, and counterstained with DAPI before observation.

#### Blockade of Reactive Astrocytes and C3/C3aR pathway in SCI Mice

To block reactive astrocytes after SCI using neutralizing antibodies (Liddelow *et al*., 2017), C57BL/6 mice rather than SD rats were selected to conduct T10 transection experiment due to the great demand of neutralizing antibodies in vivo. Expect for the Sham group (without SCI; n=12), all mice were underwent complete T10 transection SCI and injected with triple neutralizing antibodies (Neutralizing Abs group; TNFα/IL-1α/C1q, 10 mg/kg each; n=12), the control IgG antibody (IgG group; 10 mg/kg; n=12) and 0.9% normal saline (Lesion control group; n=12) via intraperitoneal injection every two days. To block the C3/C3aR pathway, C3aR antagonist (SB290157; 10 mg/kg) or 0.9% normal saline were intraperitoneally injected in SCI or Sham mice daily. And animals were treated with EdU (50 mg/kg) to track the proliferated NSCs. At seven days post-lesion, all animals were sacrificed to evaluate the NSC activation.

#### Behavior Test

The Basso, Beattie & Bresnahan (BBB) locomotor rating test (Basso *et al*., 1995) was performed at two days before injury, 1∼3 days and weekly post-surgery to evaluate the motor functions of hindlimbs. Rats were placed on a quiet and open plan to ensure spontaneous movement, then the walking and physical activities of hind limbs were observed and recorded. BBB scale comprises three parts: I. the joints movement of hind limbs, which scored 0∼7; II. the gait and coordination function of the hind limbs, which scored 8∼13; III. the fineness of the paw in motion, which scored 14∼21. Behavior evaluation was conducted weekly post-SCI by two investigators familiar with the BBB criteria but blinded to the experimental groups, separately.

#### Electrophysiology Examination

Electrophysiological evaluation was carried out as previously described (Anderson et al., 2018; Lu *et al*., 2012) at 8 weeks after SCI in rats or 6 weeks post-injury in mice. Animals were re-anesthetized and laminectomy was performed to expose the T7 and T13 spinal cord, which are both 3 segments rostral and caudal to the lesion site. A bipolar stimulating electrode spacing 2 mm was positioned at the intraspinal rostral T7 segment, and a silver ball electrode placed at the caudal T13 segment to record evoked response. 0.1 ms square wave pulse was delivered at 10 ms interval and field activity was amplified and recorded. At least 20 trails from each recording site were averaged and the amplitude of evoked potential was quantified.

#### Protein Array of Cerebrospinal Fluid from SCI Animals

Cerebrospinal fluid (CSF) was extracted using paracentesis from the cerebellomedullary cistern. Briefly, at 6 h after T10 complete transection SCI, rats (n=4) were re-anesthetized and fixed with a stereotaxic apparatus. Then the occipital protuberance was exposed, and a glass capillary was inserted into the cerebellomedullary cistern carefully through the occipital bone. Approximately 100∼120 µl CSF was collected from each rat and kept at −80°C for further investigation. CSF samples from an equal number of rats (n=4) without surgery were used as the control.

Protein microarray analyses of CSF harvested from the normal/SCI rats were performed by G-Series Rat Cytokine Array (RayBiotech, Inc., Guangzhou, China) accordingly to the manufacturers’ instructions. A total of 100 µl sample from each rat was used and raw data obtained was conducted background subtraction and normalization before clustering analysis. Gene ontology (GO) annotation, consisting of three parts including molecular function, biological process and cellular component, was applied to identify functions of potential genes. Possible signaling pathways involved in were analyzed by the Kyoto Encyclopedia of Genes and Genomes (KEGG) database.

#### Tissue Preparation

At the corresponding of experimental endpoints, the animals were perfused with 0.9% normal saline followed by 4% PFA in 0.1 M PBS (pH 7.2), then about 2∼3cm spinal tissues containing the lesion center was dissected and extracted, post-fixed for 6∼8 hours and dehydrated in 20% and 30% sucrose for 3 days successively. After embedded in Tissue-Tek O.C.T. (Sakura), 14 μm thickness of the cross sections and 1.5 cm longitudinal slices of spinal cords containing the lesion site were collected on a cryostat.

#### Immunofluorescent Staining

For immunocytochemistry analysis, the cells were fixed with 4% PFA and permeabilized using 0.5% Triton X-100, then blocked in 5% BSA for 1 hour at RT. For immunohistochemistry, the frozen sections of the spinal cord were washed in PBS for 15 mins, subsequently underwent antigen retrieval using Proteinase K and blocking with 5% BSA supplemented with 0.3% Triton X-100 for 2 hours. After the blocking procedure, the fixed cells and slices were incubated with the first antibodies overnight at 4°C. The first antibodies used in the present study included: UCHL1 (1:300; Cell Signaling Technology, CST), Nestin (1:300; Cell Signaling Technology, CST), SOX2 (1:300; Abcam), Glial Fibrillary Acidic Protein (GFAP; 1:300; Cell Signaling Technology, CST), Microtubule Associated Protein 2 (MAP2; 1:300; Abcam), Neurofilament 200 (NF200; 1:300; Cell Signaling Technology, CST), CNPase (1:300; Cell Signaling Technology, CST), NeuN (1:300; Abcam), C3a (1:10; Abcam), C3aR (1:200; Santa Cruz), NG2 (1:200; Santa Cruz), Doublecortin (DCX; 1:300; Abcam), Ki-67 (1:800; Abcam), Tubulin β3 (1:300; Cell Signaling Technology, CST), BrdU (1:1000, Sigma). Post washing in PBS for three times, cells and sections were subsequently incubated with Alexa Flour 488/594/647-conjugated secondary antibodies and counterstained with DAPI (1:5000; Sigma). All immunofluorescence images were acquired using a Zeiss LSM 880 confocal microscope.

#### Quantitative Real-time Polymerase Chain Reaction (qPCR)

Total RNA from NSCs or spinal cord tissues was extracted using Trizol (Invitrogen, life technologies) according to the reagent specification. The quantality of RNA yield was determined by the Nanodrop one (ThermoFisher). Total RNA was subjected to reverse transcription into cDNA using the PrimeScript RT reagent Kit (Vazyme). qPCR assay was conducted using the SYBR Premix EX Taq (Vazyme). The expression levels were normalized to those of glyceraldehyde 3-phosphate dehydrogenase (GAPDH). Relative expression of genes was calculated as fold changes using the 2-△△ Ct method. The primer sequences for genes are as follows. *Uchl1,* Forward primer (5’-3’): TGAAGCAGACCATCGGGAAC, Reverse primer (5’-3’): GAGTCATGGGCTGCCTGAAT; *Uchl3,* Forward primer (5’-3’): GGTCAGACTGAGGCACCAAG, Reverse primer (5’-3’): CTCATCAGGGTCGCGCTC; *Uchl5,* Forward primer (5’-3’): AGACCTTAGCAGAACACCAGC, Reverse primer (5’-3’): CAGCAGTGTACACATGTCCAAAT; *Gapdh*, Forward primer (5’-3’): TGATTCTACCCACGGCAAGTT, Reverse primer (5’-3’): TGATGGGTTTCCCATTGATGA.

#### Western Blotting

Western blot was conducted to detect the protein enrichment in cells and spinal tissues. In brief, cells and about 1 cm spinal tissues containing the lesion site were lysed in PIPA (Solarbio) supplemented with PMSF (1:100; Solarbio) and ultrasonic splitting on ice. Total of 20 µg protein was run on 10%-12% sodium dodecyl sulfate-polyacrylamide electrophoresis (SDS-PAGE) gel, transferred to polyvinylidene difluoride (PVDF; Millipore, Mississauga, Canada) membranes and blocked with 5% no-fat milk. Then membranes were incubated with first antibodies overnight at 4°C with gentle shaking, followed by combined with horseradish peroxidase (HRP)-conjugated secondary antibodies (1:1,000). Protein enrichment was visualized via chemiluminescence reagent and the grayscale analysis of bands was determined using Image J. First antibodies applied in this study are as follows: UCHL1 (1:1000; Cell Signaling Technology, CST), proteasome 20s (1:1000; Affinity Biosciences LTD), Ubiquitin (1:1000; Abcam), Nestin (1:1000; Cell Signaling Technology, CST), GFAP (1:1000; Cell Signaling Technology, CST), MAP2 (1:1000; Abcam), NF200 (1:1000; Cell Signaling Technology, CST), C3a (1:1000; Abcam), C3aR (1:500; Santa Cruz), DYKDDDDK Tag (1:1000; Cell Signaling Technology, CST), GAPDH (1:1000, Beyotime Biotechnology), β-actin (1:1000, Beyotime Biotechnology). GAPDH and β-actin were used as the internal controls.

### QUANTIFICATION AND STATISTICAL ANALYSIS

#### Quantification of Confocal Analysis

All confocal images from cells and spinal tissues were examined using a Zeiss LSM 880 confocal microscope and quantified in a blinded manner as follows. 4∼5 spinal cord sections from different levels, including the dorsal, the middle and the ventral, were selected and imaged in each rat/mouse. The sections used for quantification were selected from the same level among groups as much as possible under different conditions. The images for quantification were all captured from the lesion site of SCI animals. The diagrams that illustrate where micrographs are imaged from and the quantitative sites in related figures were shown in Figure 5A. Each data point represents once repeated experiment or acquired from one rat/mouse. All experiments were conducted at least three times biological replicates.

To quantify UCHL1-positive cells in the lesion site of spinal cord, the percentage of DAPI signals surrounded by UCHL1 was counted. To quantify the C3^+^ reactive astrocytes, SOX2^+^ NSCs, proliferated NSCs and neurons in vitro and in vivo, C3/GFAP^+^, Ki-67/Nestin^+^, SOX2/Nestin^+^, BrdU/Nestin^+^, BrdU/Tubulin β3^+^ double-staining cells were counted manually in the same field of view and the ratio of positive cells were quantified. To quantify the GFP^+^ cells infected by lentiviral vector encoding UCHL1 in the lesion center in vivo, double-staining of GFP/Nestin^+^ NSCs, GFP/NG2^+^ OPCs, GFP/Tubulin β3^+^ neurons, GFP/NeuN^+^ neurons, and GFP/GFAP^+^ astrocytes was manually counted, and the ratio of individual cell to that of the total GFP^+^ cells was calculated, respectively. To quantify the immunoreactivity of Nestin and NF-200, the relative fluorescent area of Nestin/NF-200 in the lesion center of the individual optical section was measured using Image J.

#### Statistics

SPSS (Version 20.0; Abbott Laboratories, Chicago, IL) was applied for statistical analysis. Independent, Two-tailed unpaired Student’s t-test was used for comparisons between two groups. For multiple comparisons, One-way Analysis of variance (ANOVA) followed by Tukey HSD post-hoc analysis or Kruskal Wallis test with Bonferroni correction analysis was selected. And Two-way ANOVA with Tukey HSD was conducted for BBB scores analysis. All data was presented as mean ± the standard error of the mean (SEM) and *p*<0.05 was deemed statistically significant

