## supplemental information for "UCHL1 facilitates aggregates clearance and enhances neural stem cell activation in spinal cord injury"

**ABSTRACT**

Activation of endogenous neural stem cells (NSCs) is critically important for the adult neurogenesis. However, NSC activation is extremely limited after spinal cord injury (SCI). Recent evidence suggests that accumulation of protein aggregates impedes quiescent NSC activation. Here, we found ubiquitin c-terminal hydrolase l-1 (UCHL1), an important deubiquitinating enzyme, functioned to facilitate NSC activation by clearing protein aggregations through ubiquitin-proteasome approach. Upregulation of UCHL1 enhanced NSC proliferation in the spinal cord after injury. Based on protein microarray analysis of SCI cerebrospinal fluid, it is further revealed that C3^+^ neurotoxic reactive astrocytes negatively regulated UCHL1 and aggresome clearance through C3/C3aR signaling, resulting in reduced capacity of NSC to activate. Furthermore, blockade of reactive astrocytes or C3/C3aR pathway led to enhanced NSC activation post-SCI. Together, this study elucidated a mechanism regulating NSC activation in the adult spinal cord involving the UCHL1-proteasome approach, which may provide potential molecular targets for NSC fate regulation.

**Keywords**

Ubiquitin c-terminal hydrolase l-1, protein aggregates clearance, neural stem cell activation, reactive astrocyte, complement component 3, proteasome, spinal cord injury

**This file includes:**

**Supplementary Figures 1 to 7**

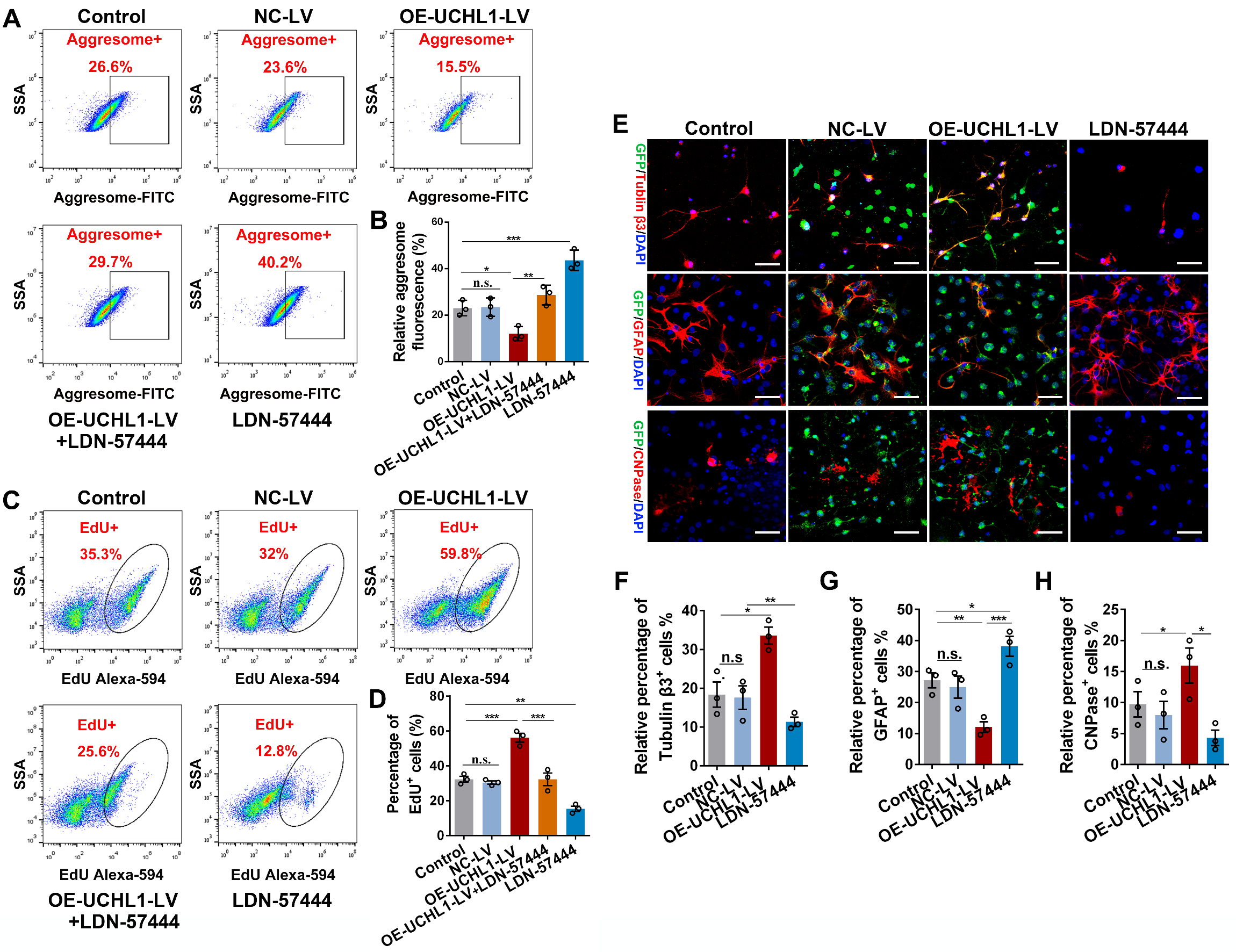

**Supplementary Figure 1. UCHL1 upregulation decreased protein aggregation accumulation, facilitated NSC proliferation and neuronal differentiation in vitro (related to Figure 2).**

(A-B) Flow cytometry analysis (A) and quantiﬁcation (B) of protein aggregates (aggresome^+^) in NSCs. (B) n=3 biological replicates.

(C-D) The proliferating NSCs (EdU^+^) in different treatments were detected by flow cytometry assay (C) and quantified (D). (D) n=3 biological replicates.

(E) Representative images showing the differentiation of NSCs treated with GFP-NC-LV, GFP-OE-UCHL1-LV or LDN-57444 for 7 days. Scale bar, 100 μm.

(F-H) Quantification of the relative ratio of differentiated neurons (Tubulin β3^+^), astrocytes (GFAP^+^) and oligodendrocyte (CNPase^+^). n=3 biological replicates.

(B/D/F/G/H) Data are presented as mean ± SEM. p-values (*p<0.05, **p<0.01, ***p<0.001, n.s. not significant) are calculated using one-way ANOVA with Tukey HSD post hoc test.

test.

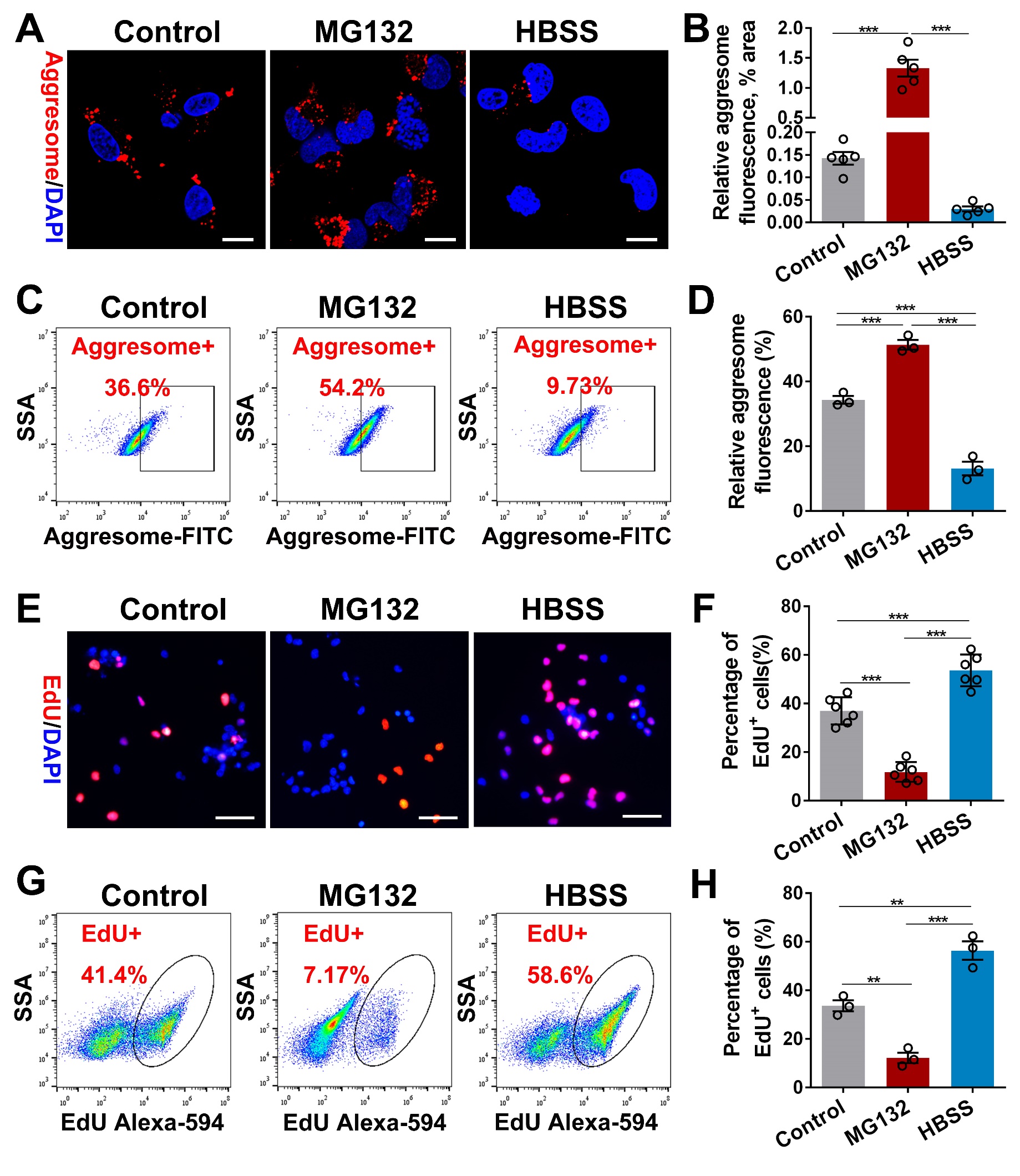
**Supplementary Figure 2. Protein aggregates accumulation in NSCs was directly related with NSC activation.**

(A) Confocal representative images showing protein aggregates labeled by aggresome dye (red) in NSCs treated with MG132 (proteasome inhibitor) or incubated with HBSS for 3 h prior to be transferred into the basal medium. Scale bar, 10 μm.

(B) Quantification of protein aggregates accumulated in NSCs. n=5 biological replicates.

(C-D) Flow cytometry analysis (C) and quantification (D) of protein aggregates in NSCs treated with MG132 or HBSS. (D) n=3 biological replicates.

(E-H) Fluorescent staining (E-F) and flow cytometry analysis (G-H) show the proliferation of NSCs treated with MG132 or HBSS for 12h. Scale bar (E), 25 μm. (F) n=6 biological replicates. (H) n=3 biological replicates.

Data are presented as mean ± SEM. p-values (**p<0.01, ***p<0.001) are calculated using (B) one-way ANOVA with Tamhane T2 post hoc test or (D/F/H) one-way ANOVA with Tukey HSD post hoc test.

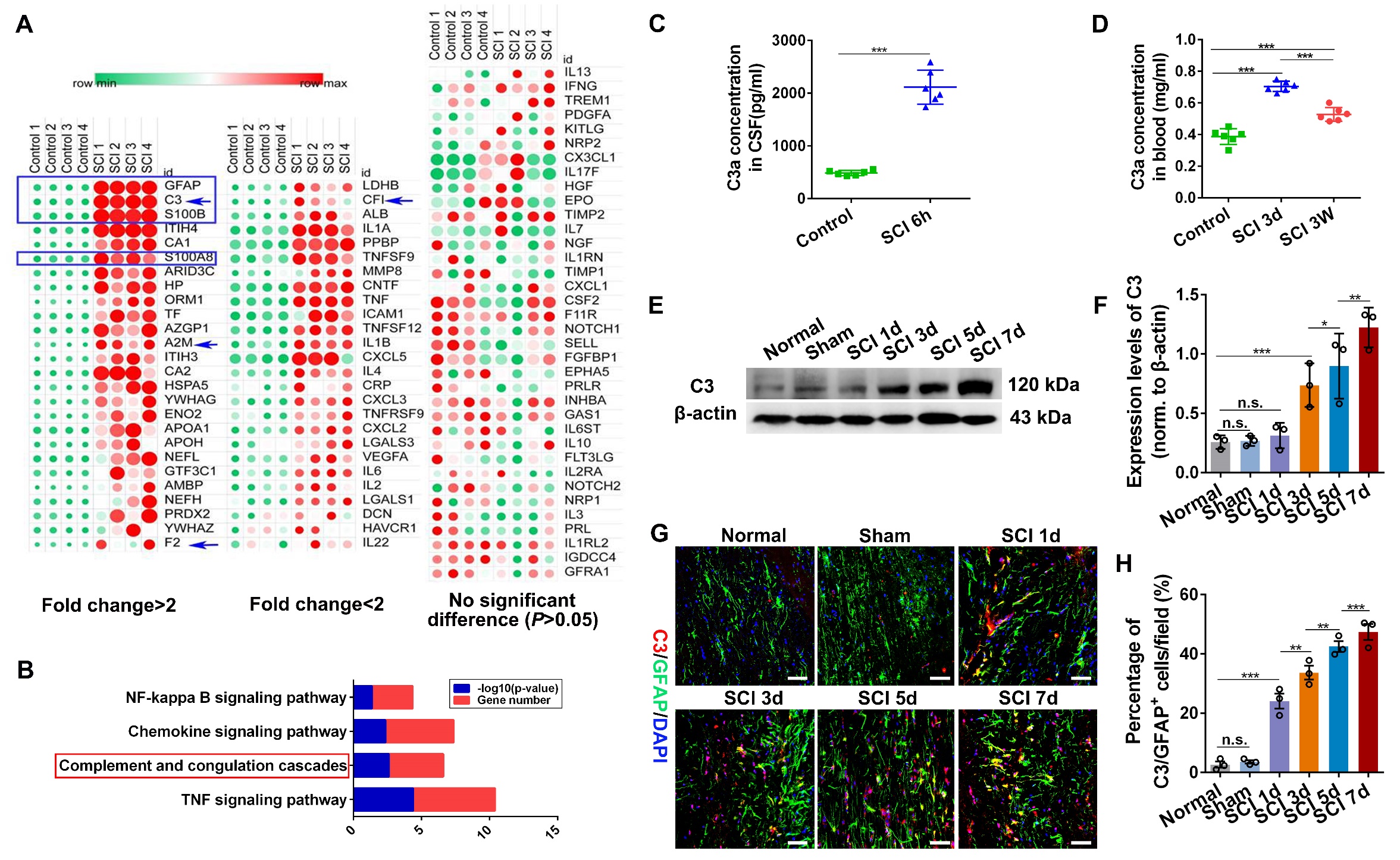

**Supplementary Figure 3. C3^+^ reactive astrocytes were abundantly activated by SCI.**

(A) Protein array of CSF from intact and SCI rats. CSF was extracted from the intact and SCI rats at 6 h post-injury and then conducted for protein microarray analysis. Scanned array image shows the relative expression changes of cytokines in CSF of SCI rats compared to that of the intact animals. The significantly changed cytokines was screened depending on conditions that the log10(fold change) ＞log10(1.2) and normalized p value <0.05. n=4 independent animals.

(B) KEGG pathway analysis of differentially expressed cytokines in CSF from intact and SCI rats. The blue bar indicates the -log transformed p value (p<0.05 is considered statistically significant), and the red bar shows the numbers of changed factors in each pathway. (C-D) The level of C3 in CSF (C) and (D) blood of SCI rats were detected by ELISA analysis. n=6 biological replicates.

(E-F) Western blot analysis and quantification of the protein level of C3 in the spinal tissues within 7 days post-SCI. (F) n=3 independent animals.

(G) Confocal representative images showing the activation of C3/GFAP^+^ astrocytes in spinal tissues within 7 days post-SCI. Scale bar, 25 μm.

(H) Quantiﬁcation of C3/GFAP^+^ reactive astrocytes in spinal tissues from Figure G. n=3 independent animals.

Data are presented as mean ± SEM. p-values (**p<0.01, ***p<0.001, n.s. not significant) are calculated using (C/D) two-tailed unpaired Student’s t-test (F/H) or one-way ANOVA with Tukey HSD post hoc test.

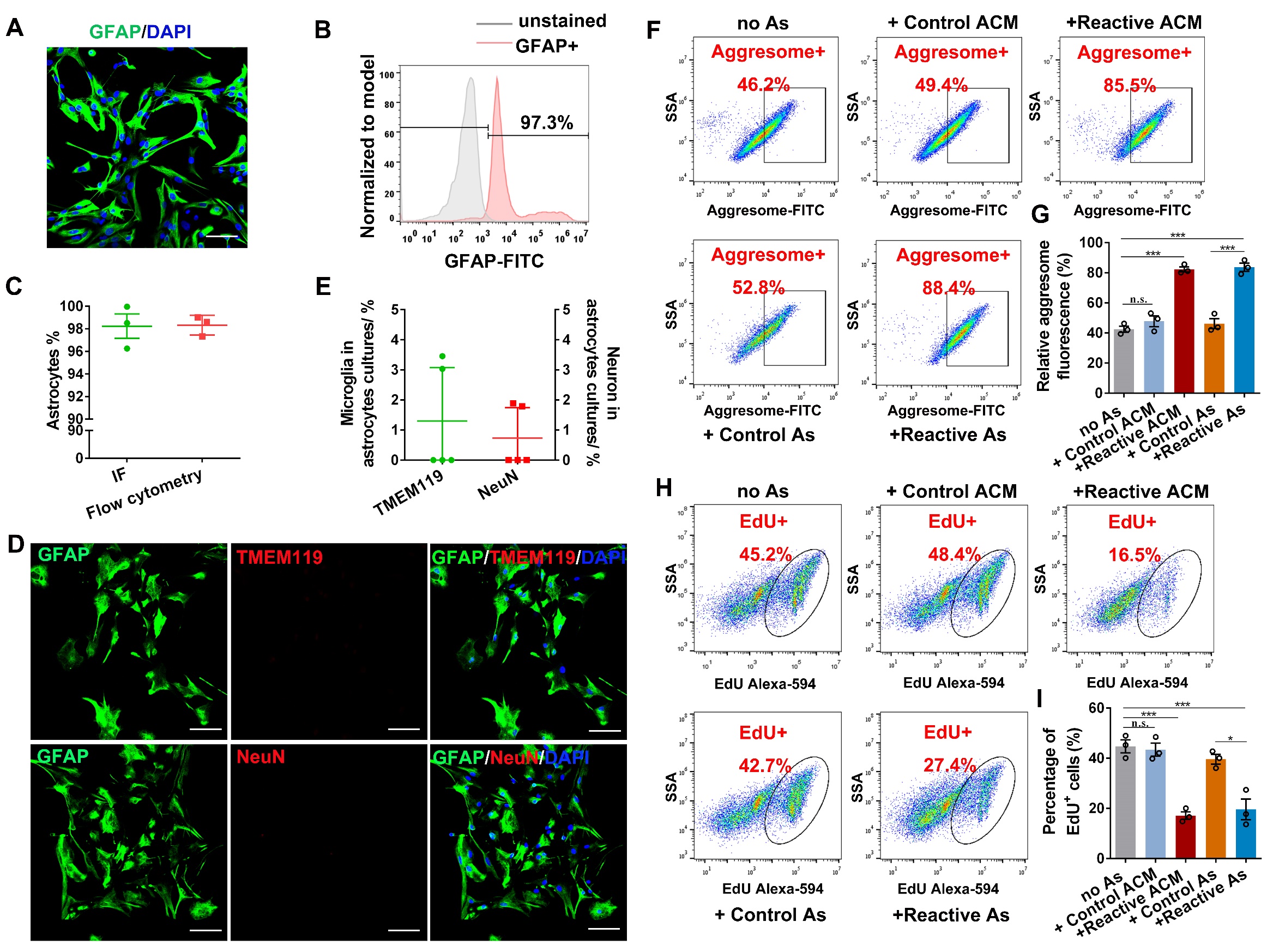

**Supplementary Figure 4. Reactive astrocytes resulted in increased protein aggregates accumulation and inhibited NSC activation by flow cytometry assay (related to Figure 3).**

(A-B) The purified primary astrocytes were confirmed by the typical marker GFAP using immunofluorescence (A) and flow cytometry analysis (B). Scale bar (A), 50 μm.

(C) Quantification of the percentage of GFAP+ cells in the primary astrocytes. The purity of astrocytes was more than 96%. n=3 biological replicates. Data are presented as mean ± SEM.

(D-E) Confocal representative images (D) showing the double staining of astrocytes (GFAP^+^) and microglia (TMEM119^+^) or neuron (NeuN^+^). Scale bar (D), 50 μm. (E) Quantiﬁcation of microglia and neuron in the primary astrocytes. n=5 biological replicates. Data are presented as mean ± SEM.

(F-G) Flow cytometry analysis (F) and quantiﬁcation (G) of accumulated protein aggregates (aggresome^+^) in NSCs co-cultured with astrocytes 24h. (G) n=3 biological replicates.

(H-I) The activation of NSCs after treated with C3 was evaluated (H) and quantified (I) by EdU^+^ NSCs by Flow cytometry assay. (I) n=3 biological replicates.

(G/I) Data are presented as mean ± SEM. p-values (*p<0.05, ***p<0.001, n.s. not significant) are calculated using one-way ANOVA with Tukey HSD post hoc test.

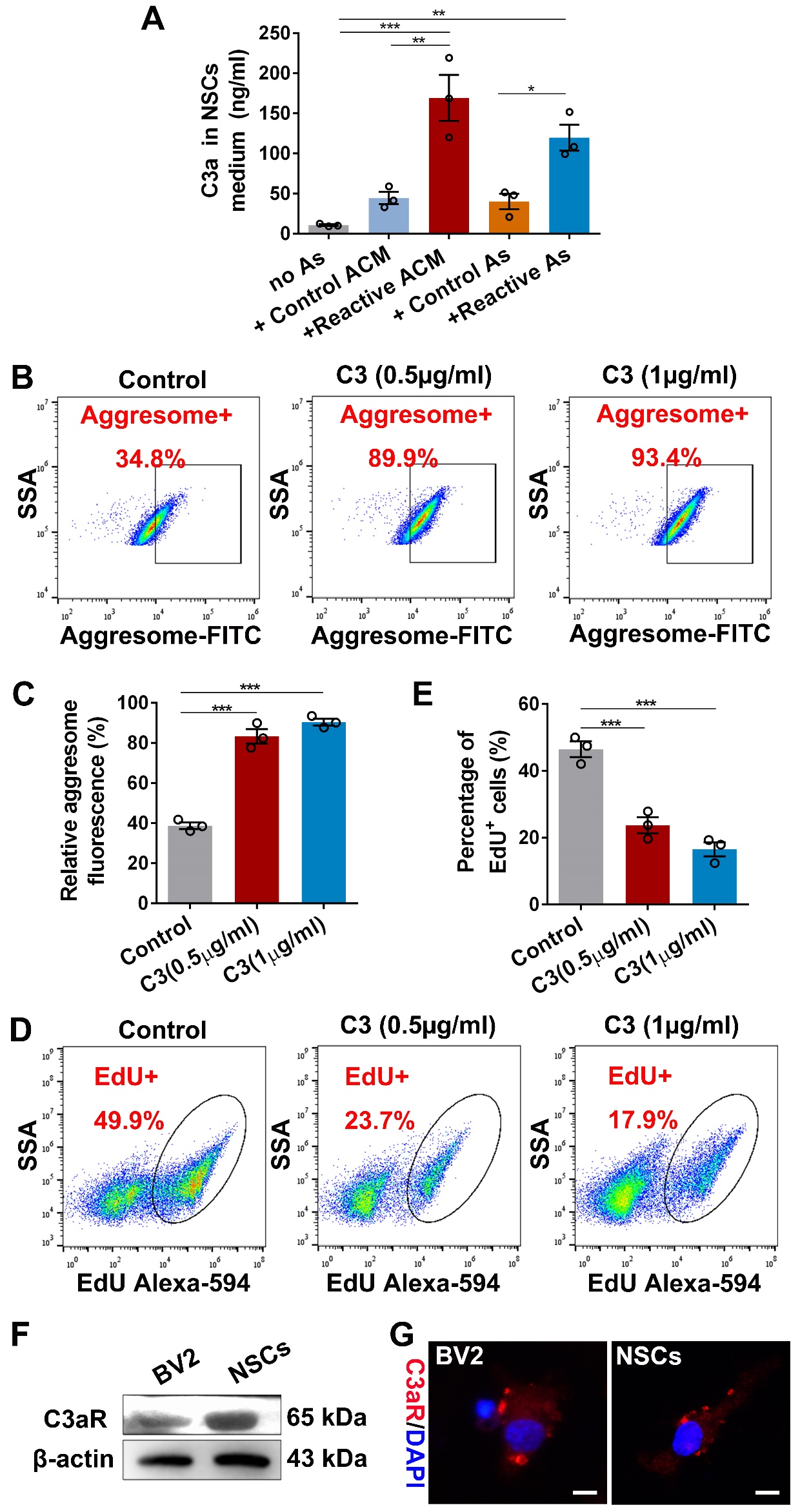

**Supplementary Figure 5. C3 enhanced protein aggregates accumulation and inhibited NSC activation by flow cytometry assay (related to Figure 4).**

(A) The level of C3a in the medium of co-culture system was measured by ELISA assay. n=3 biological replicates.

(B-C) Flow cytometry analysis (B) and quantiﬁcation (C) of accumulated protein aggregates (aggresome^+^) in NSCs treated with C3 for 24h. (C) n=3 biological replicates.

(D-E) The activation of NSCs after treated with C3 was evaluated (D) and (E) quantified by EdU^+^ NSCs through Flow cytometry analysis. (E) n=3 biological replicates.

(F-G) Western blot analysis and immunofluorescence assay show the expression of C3aR in BV2 and NSCs. Scale bar (G), 10 μm.

(A/C/E) Data are presented as mean ± SEM. p-values ((*p<0.05, **p<0.01, ***p<0.001) are calculated using one-way ANOVA with Tukey HSD post hoc test.

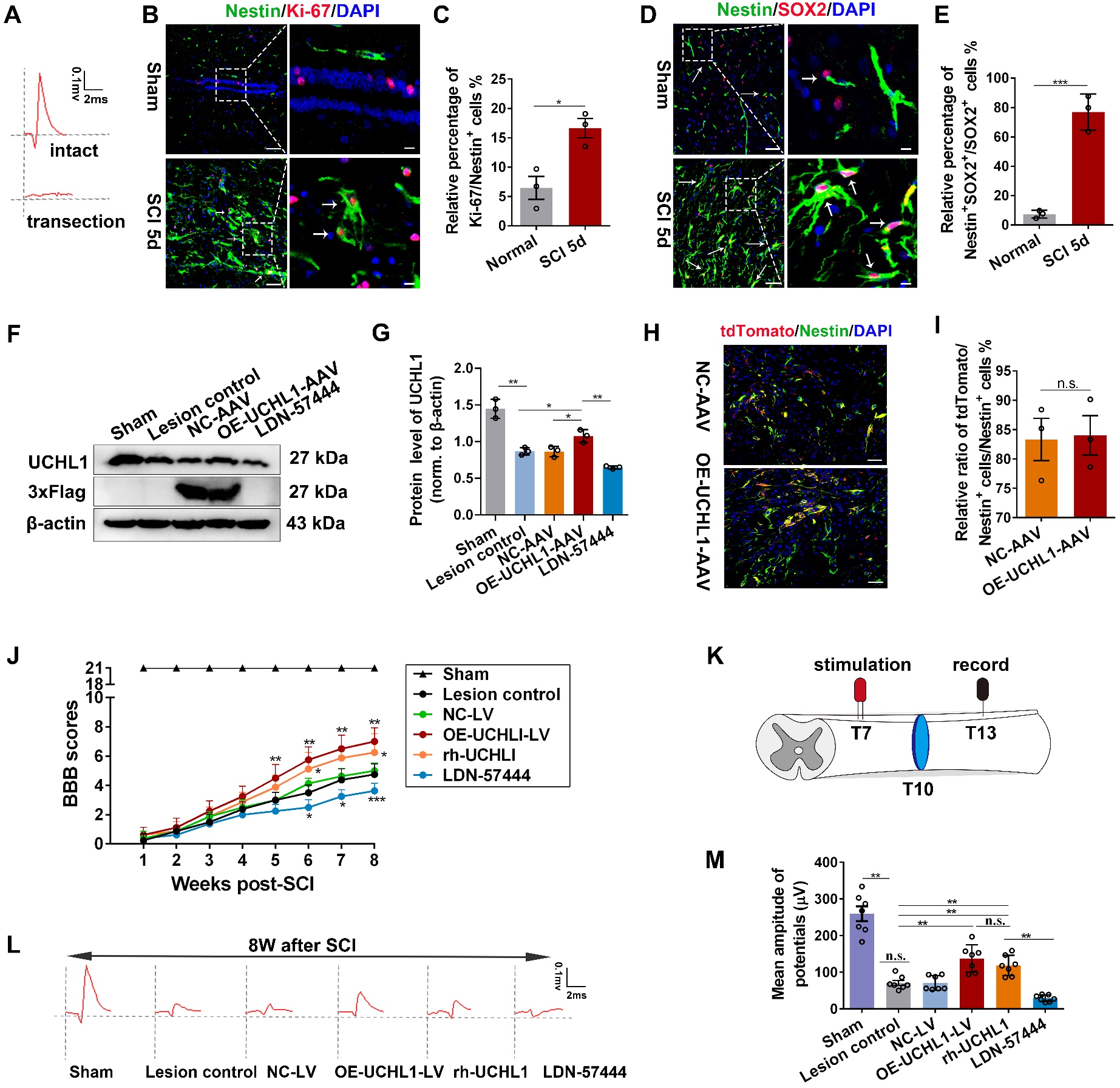
**Supplementary Figure 6. Overexpression of UCHL1 was achieved by AAV**

**encoding UCHL1 in vivo and UCHL1 upregulation improved locomotor function recovery after SCI.**

(A) Electrophysiological assay was conducted to confirm the complete transection SCI model.

(B) Validation of the proliferation ability of Nestin^+^ cells at 5-day after SCI. Scale bar, 25 μm.

(C) Quantification of Ki-67/Nestin^+^ cells at 5 days post-SCI. n=3 independent animals. Data are presented as mean ± SEM. p-values (*p<0.05) is calculated using two-tailed unpaired Student’s t-test.

(D) Activation of Nestin/SOX2^+^ cells at the acute phase of SCI rats. Nestin/SOX2^+^ cells were largely activated around the center lesion post-SCI. Scale bar, 20 μm.

(E) The relative ratio of Nestin/SOX2^+^ cells to that of SOX2+ cells were quantified. n=3 independent animals. Data are presented as mean ± SEM. p-values is calculated using two-tailed unpaired Student’s t-test.

(F-G) Western blot analysis of spinal tissues lysate from SCI rats treated with OE-UCHL1-AVV or LDN-57444 at two weeks post-injury. Quantification of the relative expression of UCHL1 in spinal tissues was shown in I. n=3 independent animals. Data are presented as mean ± SEM. p-values (*p<0.05, **p<0.01) are calculated using one-way ANOVA with Tukey HSD post hoc test.

(H-I) Confocal representative images and quantification of tdTomato/Nestin^+^ NSCs in spinal at 2 weeks after SCI among different treatments. Scare bar (J), 25 µm. n=3 independent animals. Data are presented as mean ± SEM. p-values (no significance) is calculated using two-tailed unpaired Student’s t-test.

(J) BBB scores of SCI rats administrated with OE-UCHL1-LV, rh-UCHL1 or LDN-57444 during the 8 weeks post-injury. n=8 independent animals. Data are presented as mean ± SEM. p-values (* vs Lesion control: *p<0.05, **p<0.01, ***p<0.001) are calculated using Two-way ANOVA.

(K) Schematic diagram of electrophysiological assay.

(L) Rats were subjected to electrophysiological assay at 8 weeks before perfusion to assess the reestablishment of neural circuit. Representative images of the provoked response were recorded and shown in C.

(M) The amplitude of evoked potential was quantified by the peak-to-peak value. n= 7 independent animals. Data are presented as mean ± SEM. p-values (**p<0.01, n.s. not significant) are calculated using one-way ANOVA with Kruskal Wallis test with Bonferroni correction.

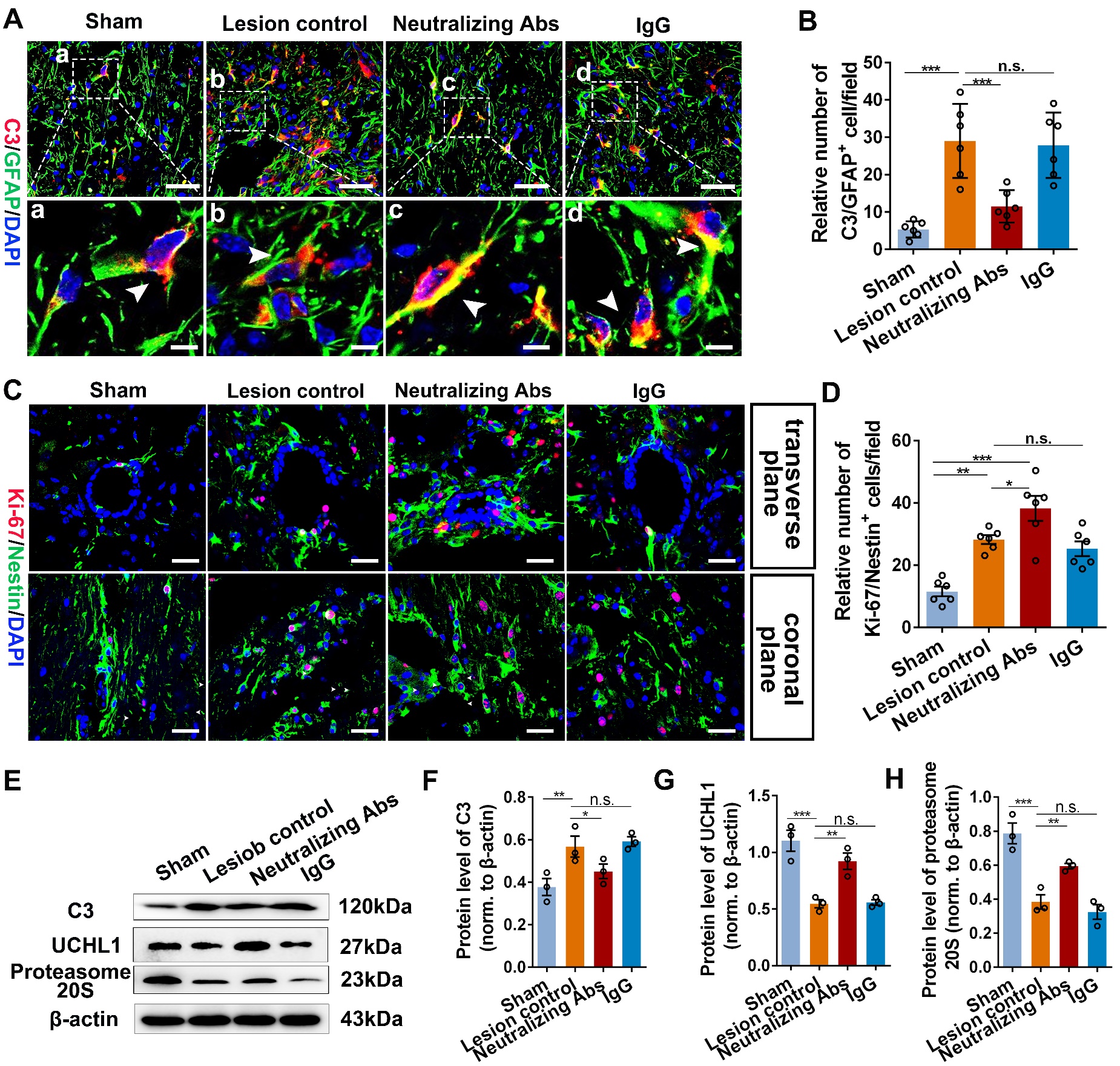

**Supplementary Figure 7. Blockade of reactive astrocytes using neutralizing antibodies effectively enhanced NSC proliferation in SCI mice.**

(A) Representative images showing the activation of reactive astrocytes (C3/GFAP^+^) detected by immunofluorescence assay at 7 days post-SCI. Administration of neutralizing antibodies significantly blocked formation of reactive astrocytes after SCI. Enlarged images of the boxed region are shown in the bottom panels. Scale bar (A), 20 μm. Scale bar (a-d), 10 μm.

(B) Quantification of C3^+^ reactive astrocytes in the lesion site at 7 days post-SCI. n=6 independent animals.

(C) Proliferation of NSCs around the central canal (transverse plane) and in the lesion center (coronal plane) was evaluated by the detection of Ki-67^+^ NSCs via immunofluorescence assay. Scale bar, 20 μm.

(D) The relative number of Ki-67/Nestin^+^ cells per field were counted. n=6 independent animals.

(E-H) Western blotting assay of spinal lysate from SCI mice treated with neutralizing antibodies or IgG isotype control. Representative immunoblot images and quantification of the relative enrichment of proteins were revealed in E and F/G/H. (F/G/H) n=3 independent animals.

(B/D/F/G/H) Data are presented as mean ± SEM. p-values (*p<0.05, **p<0.01, ***p<0.001, n.s. no significant) are calculated using one-way ANOVA with Tukey HSD post hoc test.
